# The interaction of compliance and activation on the force-length operating range and force generating capacity of skeletal muscle

**DOI:** 10.1101/580027

**Authors:** S.M. Cox, K.L. Easton, M. Cromie Lear, R.L. Marsh, S.L. Delp, J. Rubenson

## Abstract

Muscle performance is influenced by where it operates on its force-length curve. Here we explore how activation and tendon compliance interact to influence muscle operating lengths and force-generating capacity. To study this, we built a musculoskeletal model of the lower limb of the guinea fowl and simulated the force-length operating range during fixed-end fixed-posture contractions for 39 actuators under thousands of combinations of activation and posture using three different muscle models: Muscles with non-compliant tendons, muscles with compliant tendons but no activation dependent shift in optimal fiber length (L0), and muscles with both compliant tendons and activation-dependent shifts in L0. We found that activation dependent effects altered muscle fiber lengths up to 40% and increased or decreased force capacity by up to 50% during fixed-end contractions. Typically, activation-compliance effects reduce muscle force and are dominated by the effects of tendon compliance at high activations. At low activation, however, activation-dependent shifts in L0 are equally important and can result in relative force changes for low compliance muscles of up to 60%. There are regions of the force-length curve in which muscles are most sensitive to compliance and there are troughs of influence where these factors have little effect. These regions are hard to predict, though, because the magnitude and location of these areas of high and low sensitivity shift with compliance level. Here we provide a map for when these effects will meaningfully influence force capacity and an example of their contributions to force production during a static task, namely standing.

## Introduction

Where muscles operate on their force-length relationship has important implications for muscle and locomotor performance. Most tangibly, muscle length affects muscle force and, therefore, joint torque capacity (Arnold et al., 2013; Blix, 1894; Gordon et al., 1966; Huxley and Simmons, 1971; Morgan et al., 2002; Rack and Westbury, 1969; Ramsey, R. W. & Street, 1941; Rassier et al., 1999; TerKeurs et al., 1978). One relatively unexplored, but potentially significant, factor influencing muscle operating lengths is muscle–tendon compliance. Zajac (Zajac, 1989) initially suggested that even in “isometric” (fixed-end) contractions, muscle fibers will shorten due to the stretch of the tendon, an effect exaggerated at higher activation levels due to greater forces. The result is that the operating ranges for muscles with compliant tendons shift to the left on the force-length curve with increasing activation. This theoretical prediction has been experimentally confirmed in several studies (Arnold and Delp, 2011; Azizi and Roberts, 2010; Fukunaga et al., 1997; Hawkins and Bey, 1997; Holt and Azizi, 2016; Lakatta and Jewell, 1977; Lemos et al., 2008; MacIntosh and MacNaughton, 2005; Mayfield et al., 2016; Rubenson et al., 2012; Sugisaki et al., 2011). More recently, it has been shown that tendon compliance is an important factor affecting muscle force-length operating ranges during human gait (Arnold and Delp, 2011). Yet, despite many studies demonstrating this effect for individual muscles or during specific tasks, we still do not have a broad understanding of when tendon compliance has a meaningful influence on muscle operating length and function. For instance: Is there a threshold of muscle-tendon compliance where the effect becomes functionally relevant? Does compliance have a consistent influence on the F-L operating range in all conditions (i.e. across different postures and muscle lengths) or are there conditions when it is more influential?

Complicating these questions is the observation that, even in the absence of muscle-tendon compliance, activation can alter optimal muscle lengths. While the exact mechanisms remain unclear, a rightward shift of the plateau region of the force-length curve occurs with decreasing activation (Holt and Azizi, 2014; Rack and Westbury, 1969; Roszek et al., 1994) such that the optimal muscle length for force production increases. Although an interaction between activation and compliance on muscle operating lengths has been demonstrated (Arnold and Delp, 2011; Arnold et al., 2013; Lemos et al., 2008; Lieber and Brown, 1992; Rubenson et al., 2013), no studies have systematically explored the simultaneous influence of activation-dependent optimal fiber length and tendon compliance, and its subsequent effect on force-length operating ranges. Nor have the relative contributions of compliance and activation dependent shifts of the force-length curve been quantified. Therefore, we do not know under what conditions we can or cannot safely ignore these complicating factors in interpreting and predicting how muscle force varies with length across any species.

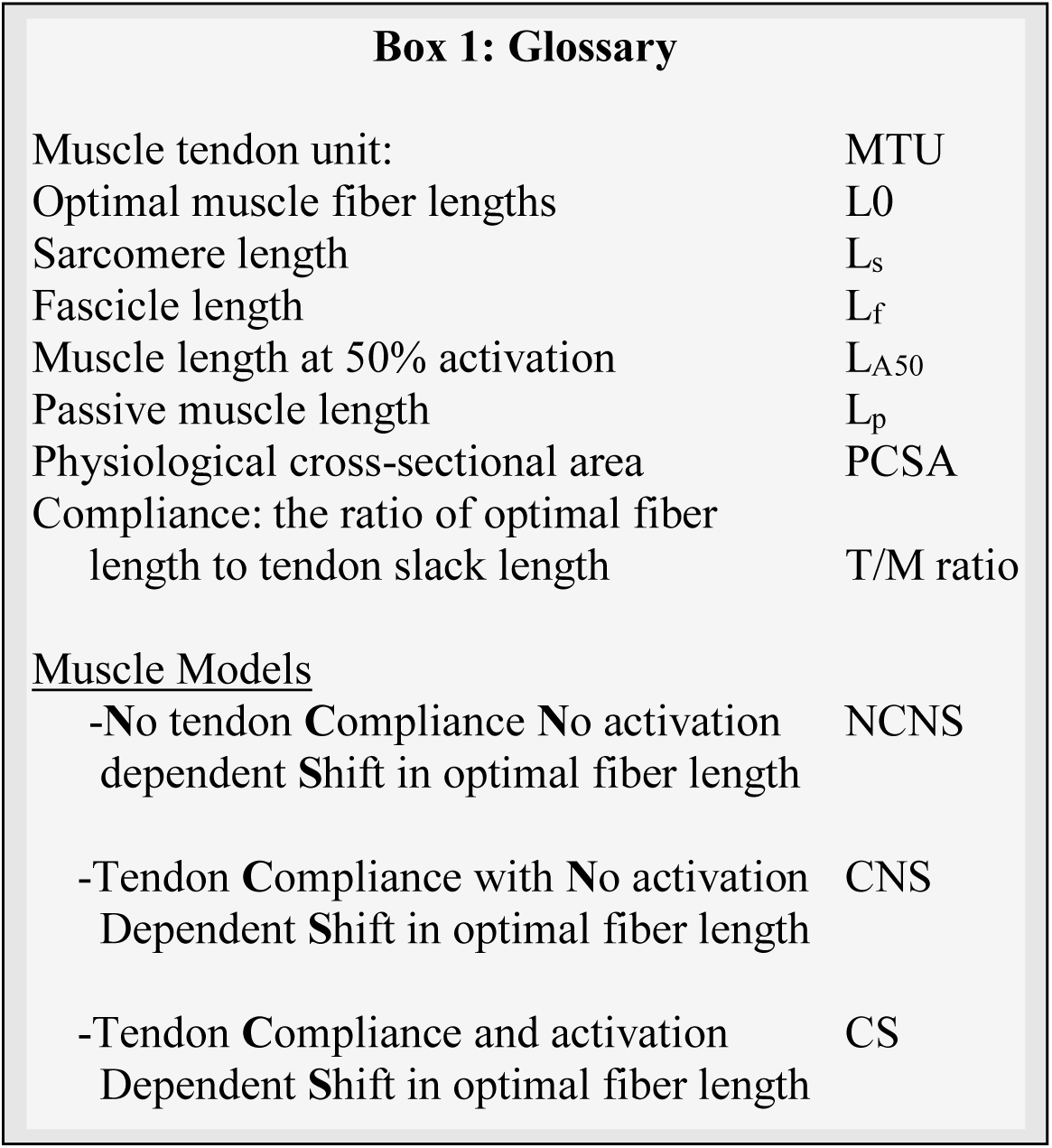

The purpose of this study is to undertake the first comprehensive assessment of how tendon elasticity and activation-dependent shifts in optimal fiber lengths combine to influence force-length operating ranges. To this end, we integrated a comparative and computational approach, the combination of which is well suited to illuminate these relationships. We developed a computational musculoskeletal model of the guinea fowl pelvic-limb (Fig. 1A), a popular avian bipedal model for biomechanical studies (Daley and Biewener, 2011; Daley et al., 2006; Ellerby et al., 2005; Ellerby and Marsh, 2010; Gordon et al., 2015; Marsh et al., 2006; Rubenson and Marsh, 2009). An animal computational model has the advantage of including highly detailed measurements of musculoskeletal parameters, many of which are typically not included in current human models (see Supplemental Material). Additionally, in this animal model the scope of muscle-tendon compliance (Table 1) is twice that of humans (Arnold and Delp, 2011), amplifying the effects. Using a computational approach allowed us iterate many more combinations of activation and posture than would be possible experimentally. We simulated the force-length operating range for 39 actuators (Table 1) under thousands of combinations of activation and posture to distill the influence of tendon elasticity and activation on force-length operating range (Fig. 1). This computational approach enabled us to extrapolate overall patterns and to systematically tease out the contributions of individual factors; for example, the role of the initial passive (pre-activation) lengths of muscles, activation dependent optimal fiber lengths, and tendon compliance on muscle operating length.

**Figure 1.**
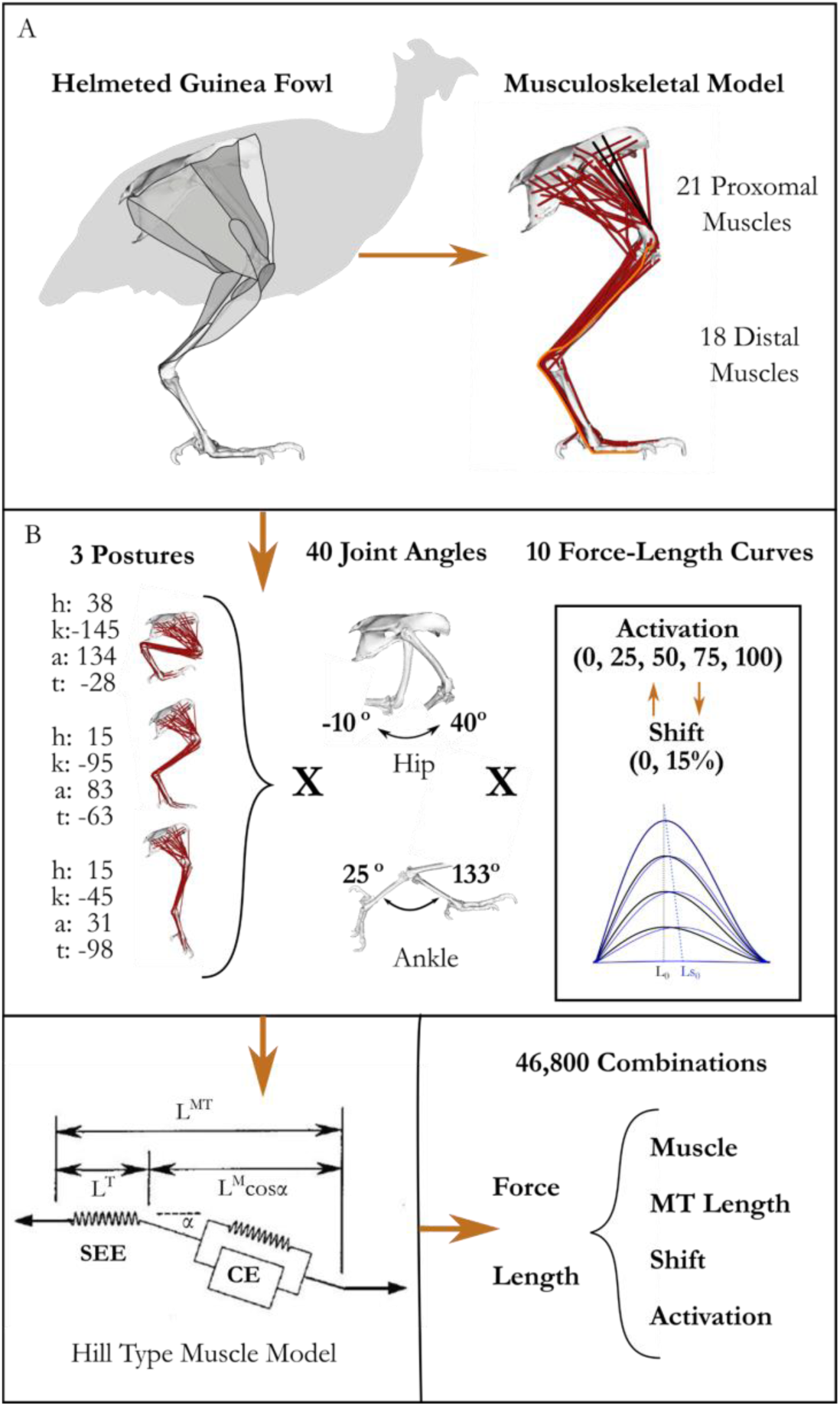
Methodological approach. A) Musculoskeletal model developed in SIMM and implemented OpenSim (see supplemental material. B) Since several muscles cross multiple joints, we varied starting MTU length by changing the posture and initial joint angle.

**Table 1.** Tendon slack length (TSL), optimal fiber length (OFL) and their ratio listed for each muscle used for analysis. Muscles are organized by group (proximal vs distal) and mean T/M Ratio (TSL/OFL) for each group listed. For full muscle names see Supplemental Material.

| Muscle | TSL(m) | OFL(m) | T/M Ratio | Muscle | TSL(m) | OFL(m) | T/M Ratio |
| --- | --- | --- | --- | --- | --- | --- | --- |
| Hip flexion/Extension Muscles |  |  |  | Ankle Flexion/Extension Muscles |  |  |  |
| CFP | 0.027 | 0.037 | 0.71 | DFii_d | 0.182 | 0.011 | 16.55 |
| FCLP_c | 0 | 0.120 | 0.00 | DFii_dx | 0.185 | 0.015 | 12.33 |
| FCLP_p | 0.078 | 0.054 | 1.46 | DFiii_dx | 0.196 | 0.016 | 12.33 |
| IC_cr | 0 | 0.086 | 0.00 | DFiii_d | 0.195 | 0.013 | 14.77 |
| IC_cd | 0 | 0.085 | 0.00 | EDL_iii | 0.193 | 0.012 | 15.69 |
| ILPO_cr | 0.064 | 0.033 | 1.93 | FHL_iii | 0.239 | 0.020 | 11.75 |
| ILPO_m | 0.017 | 0.083 | 0.21 | FL_l | 0.084 | 0.049 | 1.73 |
| ILPO_cd | 0 | 0.111 | 0.00 | FL_p | 0.081 | 0.049 | 1.67 |
| ILPR_cr | 0.044 | 0.041 | 1.09 | IG | 0.075 | 0.072 | 1.05 |
| ILPR_cd | 0.057 | 0.026 | 2.19 | LG | 0.102 | 0.047 | 2.17 |
| ISF_v | 0.039 | 0.014 | 2.76 | MG_l | 0.105 | 0.041 | 2.57 |
| ISF_d | 0.039 | 0.014 | 2.76 | MG_c | 0.118 | 0.037 | 3.17 |
| ITC_d | 0.019 | 0.023 | 0.83 | MG_m | 0.120 | 0.028 | 4.23 |
| ITC_v | 0.012 | 0.028 | 0.42 | TC_f | 0.119 | 0.029 | 4.07 |
| ITCR | 0.006 | 0.023 | 0.28 | TC_t | 0.072 | 0.051 | 1.42 |
| ITM | 0.008 | 0.023 | 0.33 | DFii_s | 0.198 | 0.013 | 15.19 |
| PIFL_cd | 0 | 0.075 | 0.00 | DFiii_s | 0.183 | 0.037 | 4.97 |
| PIFL_cr | 0 | 0.049 | 0.00 | FDL_iii | 0.214 | 0.037 | 5.78 |
| PIFM_cd | 0 | 0.075 | 0.00 | **Mean TRF** |  |  | **7.82** |
| PIFM_cr | 0 | 0.040 | 0.00 |  |  |  |  |
| FCM | 0.003 | 0.74 | 0.01 |  |  |  |  |
| **Mean TFR** |  |  | **0.71** |  |  |  |  |

We addressed three questions: 1) Averaging across all muscles, how does activation level interact with muscle-tendon unit (MTU) compliance, initial passive muscle length and/or activation-dependent shifts in L0 to alter the length and force generating capacity of muscles? 2) Under what combinations do these factors meaningfully influence force generating capacity of an individual muscle? 3) How do these factors influence the force and/or torque generating capacity during a low-activation isometric task like standing?

## Methods

### OpenSim Model development

Our approach to explore these questions was to build a realistic detailed musculoskeletal model. This was done first in SIMM software (Delp and Loan, 1995) and subsequently converted to OpenSim (Seth et al., 2018). This model incorporated several experimental measurements of muscle and skeletal properties as well as steps to validate the model accuracy (Fig. 2). A detailed description of each modelling step is provided in the Supplemental Material and summarized below.

**Figure 2.**
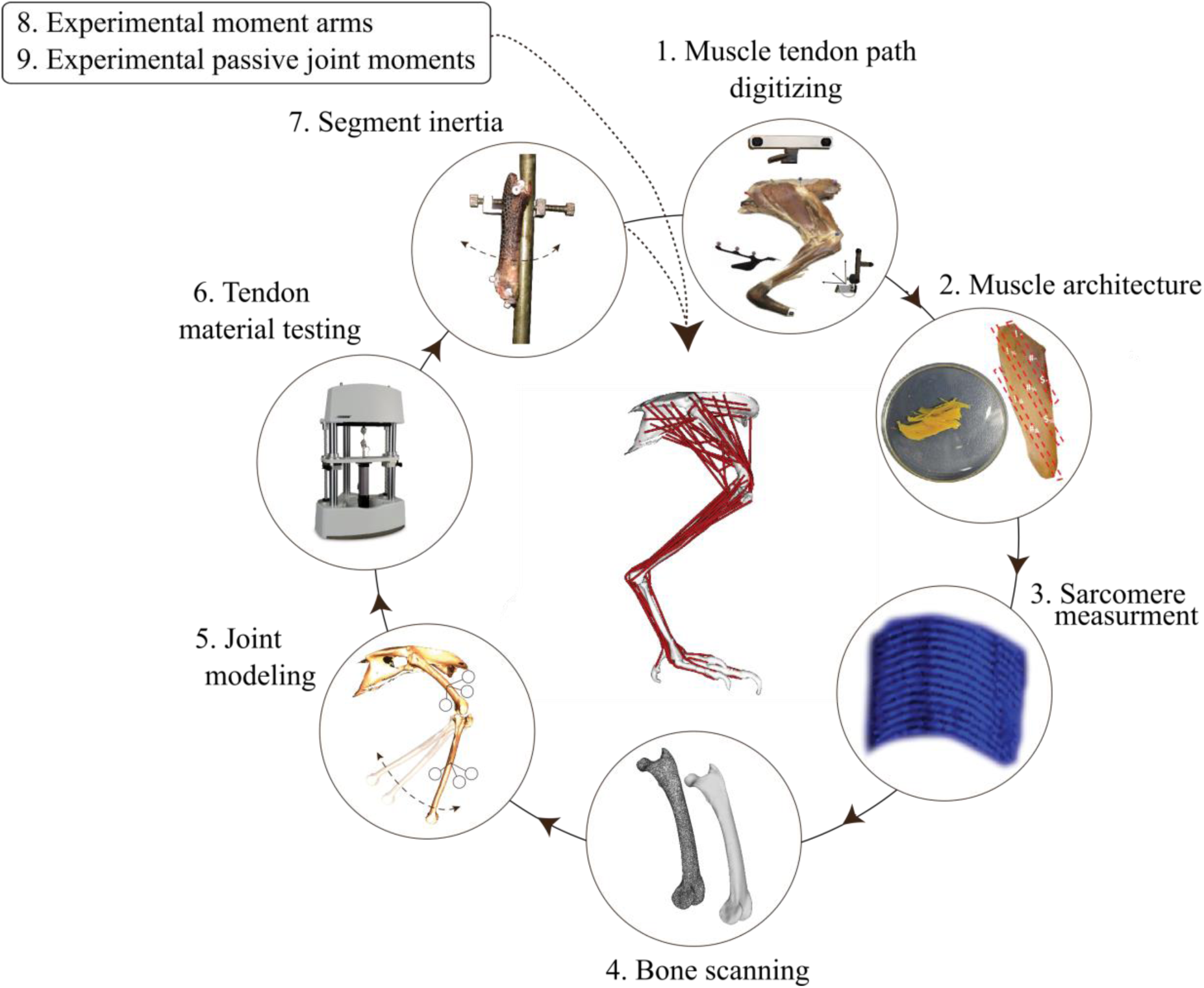
Model development framework and main steps. See Methods section and Supplementary Material for details.

The modeling began by digitizing the 3D muscle-tendon paths of the model animal (1.45 kg). This was achieved by isolating the pelvic limb and systematically dissecting off each muscle-tendon-unit (MTU) from the right limb (kept fresh/frozen). The muscle-tendon path from origin to insertion was traced using an optical tracking system (Polaris, Northern Digital, Waterloo, ON). A total of 39 MTUs were defined for this study (for a complete muscle list of the model see Supplemental Material), with some large muscles being divided into sub-muscles. Care was taken to capture the geometry of MTU paths across articulating surfaces, which were used to define muscle wrapping surfaces in SIMM software. Muscle architecture was measured on the model specimen as well as four additional animals (1.46 ± 0.1 kg; mean ± SD) for comparison. Experimental muscle architecture measurements included the muscle mass and free tendon mass and length. The left limb was fixed (10% neutral buffered formalin) in mid-swing posture and used to measure muscle fiber lengths (L_f_) and pennation angles. Bundles of fascicles were isolated and used for sarcomere length measurements (L_s_) based on second harmonic generation using two-photon laser microscopy (Cromie et al., 2013). Optimal muscle fiber lengths (L0) were defined as *L*0 = *L*_*f*_ ⋅ ^2.36^⁄_*L*_*s*__, where 2.36 is the length in microns of the optimal sarcomere length in guinea fowl muscle (Carr et al., 2011; Llewellyn et al., 2010). The Cross-sectional Area (*CSA*) of the muscle was calculated as from the L0, muscle mass (*m_mus_*) and density (*ρ*_*mus*_, 0.00106 g/mm^3^) as: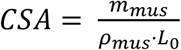. Maximal isometric force (F_100max_) for each muscle unit was calculated using a specific tension of 0.3 N/mm^2^.

Anatomical and functionally relevant bone coordinate systems (B-ACS) were defined, incorporating mathematically derived joint centers and axes. For the knee, ankle and tarsometatarsus-phalangeal (TMP) joint, a helical axis and functional joint center were computed from 3D motion capture data of adjacent segment motion (Rubenson et al., 2007). Ankle joint translation and patella-complex motion was defined as a function of ankle and knee joint angle, respectively (measured from 3D motion capture). The hip joint center was defined by directly digitizing its location with the 3D pointer. Skeletal elements were 3D-scanned to generate bone mesh (.ply) files. The bone model files and muscle-tendon paths were transformed into the relevant B-ACSs that were used to define the final model’s joint coordinate systems (JCS; see Supplemental Material for a full definition of B-ACSs and JCSs).

We measured tendon material properties from two separate animals (1.43 kg; 1.49 kg) of the free common tendon of the lateral, medial and intermedius gastrocnemius muscles (Achilles), and the free tendon from the tibialis cranialis, digital flexor IV and extensor digitorum longus (Bose EnduraTEC, ELectroForce 3200, Framingham, MA, USA). The tendons were programmed to undergo a 5Hz sinusoidal cycle that approximated the duration of the stance phase / swing phase of gait (Rubenson and Marsh, 2009). The clamp displacement was programmed to produce force approximating F_100max_. These data were used to generate muscle-specific tendon load-elongation curves for the tested muscles and to generate a generic (average) load-elongation curve for the remaining tendons. The tendon slack lengths were defined by constraining the passive simulated muscle fiber length to match the experimental fiber length in joint postures matching the experimental specimen. Segment inertial properties for the model were obtained from previously collected data from six animals (See Rubenson and Marsh, 2009).

Finally, to assess the accuracy of our model we compared simulated data to experimental values for both muscle moment arms and passive net joint moments. Muscle moment arms were computed from tendon travel experiments measured post mortem in separate specimens (see Supplemental Material). Passive net joint moments were measured for the hip joint (proximal muscles) and ankle joint (distal muscles) from deeply anesthetized and nerve-blocked animals, separate to that of the model specimen (1.55 ± 0.2 kg; mean ± SD). Details of these experiments and results are presented in the Supplemental Material.

### Computation of fiber lengths during isometric (fixed-posture) contractions

In this musculoskeletal model, we simulated fixed-end contractions across a range of activation levels and limb configurations (Fig. 1B) intended to elicit the maximum *in vivo* range of muscle lengths. MTU lengths were roughly prescribed by setting a posture that set an initial passive length (short, medium and long) and then imposing smaller deviations in MTU lengths around that initial position though variation of either the ankle (distal muscles) or hip angle (proximal muscles) through 95% of the range for that joint during locomotion reported in the literature across various experiments (Ellerby and Marsh, 2006, 2010; Henry et al., 2005; Rubenson and Marsh, 2009). Each posture generated a range of passive MTU lengths across the limb determined by fiber length, moment arm and tendon slack length of each muscle. At each of these postures we implemented three different muscles models (1: No Compliance, No activation-dependent shift in L0 (NCNS), 2: Compliance with No activation-dependent shift in L0 (CNS) and, 3: Compliance with a 15% activation dependent Shift in L0 (CS) and simulated fixed-end contractions at 5 activation levels (0%, 25%, 50%, 75%, 100%). Simulations for muscle models without activation-dependent shift were performed on the unaltered model with and without tendon compliance. Including activation dependent shifts of optimal fiber length required altering the force-length curve for activation levels below 100%. For each activation level, the force-length curve for all muscles was replaced in the OpenSim model with a shifted curve (see supplemental material for details). A 15% activation-dependent shift in L0 (at zero activation) was used because it represents a typical value reported in the literature for both voluntary muscle activation and electrically stimulated isolated muscle under realistic stimulation frequencies (de Brito Fontana and Herzog, 2016; Rack et al., 1969; Roszek et al., 1994), and also because it is a value previously adopted in modeling of muscle force (Buchanan et al., 2004; Lloyd and Besier, 2003). For each simulation, we extracted the normalized active individual fiber force and normalized fiber length for each muscle. Passive muscle lengths were defined as the muscle length at 0% activation. This resulted in 46,800 MTU lengths (Fig. 1B). These results were analyzed in three ways.

**1. On average, how does activation level interact with MTU compliance, initial passive muscle length and/or activation-dependent shift in L0 to influence the operating range and force capacity of muscles?**

To discern whether there are any useful broad patterns of interaction between MTU compliance, (i.e. the ratio of tendon slack length to optimal fiber length, Table 1), initial passive muscle length and activation dependent shifts in L0, we binned data into two levels of compliance (Low< 2< High), and passive muscle length [ascending (A), plateau (P) or descending (D) limb of the force-length curve] and evaluated how muscle length changed with activation level between muscle models. For each group (see Table 1), we calculate average normalized muscle length [L/(L0_100_)] and force (F/ F_100max_) across all muscles in each group (Table 2) at each level of activation (0%,25%,50%,75%,100%). Additionally, at each activation level we calculated how muscle length changed with activation (ΔL) from its passive starting length. For instance, ΔL at 50% activation would be given by

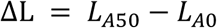

**Table 2.**
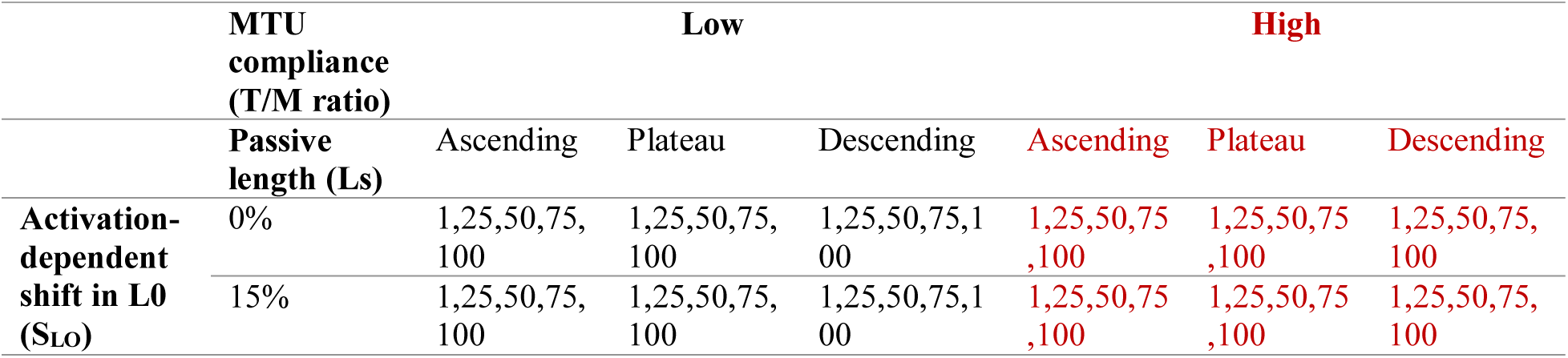
Triple nested experimental design

Where the muscle length at 50% and 0% activation (normalized to L0 at 100% activation) for this passive length is given by L_A50_ and L_A0_ respectively. Main and interaction effects were evaluated with a nested linear mixed effects model (Bates et al., 2015) with ΔL as the dependent variable, muscle as a random factor and activation level (0,25,50,75,100), activation-dependent shift in L0 (0,15 %), passive length of the muscle (A, P, D) as independent factors. Given that activation dependent shifts in the optimal fiber length did not show a main or any interaction effects, the statistical models were run both with and without this factor. Throughout this paper, muscle lengths are reported as normalized by the optimum fiber length at 100% activation, L0_100_, and forces by maximum force at 100% activation, F_100max_.

**2. Under what combinations of these factors is the force generating capacity of an individual muscle meaningfully altered?** Specifically, we asked how compliance and activation-dependent shift in L0 alter muscle operating length and force capacity at high (100%) and low (25%) activation and which combinations of these factors alter the relative force capacity substantially.

Our second analysis aimed to quantify our results at a finer level of detail. While in the first analysis we binned our data into two compliance levels and three different passive length ranges, here we present the length and force data for each muscle at all simulated passive lengths (9216 iterations). Again, we isolated the influence of compliance and activation-dependent shifts in L0 by comparing resultant muscle lengths and forces between the three muscle-tendon models (NCNS, CNS, CS). Differences between models with and without compliance isolate the influence of compliance alone. Likewise, differences between models with compliance but with and without activation-dependent shifts in L0 allow us to isolate the contribution of activation-dependent shift to changes in operating length and force capacity. Specifically, the contribution of activation-dependent shift in L0 to muscle operating length (ΔL) for a given muscle, passive length and activation level was found by subtracting the muscle length of the CNS condition, L_CNS_, from that of the CS condition, L_CS_.

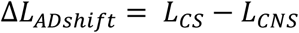

The force values for each muscle with no compliance or shift (i.e. non-compliant tendon, NCNS) were found by assuming no change in muscle length across activation levels and scaling the force with activation. For example, for a muscle with a passive length of 0.809 at 100% activation, F_100_, is 0.726. At 50% activation, the calculated force with no compliance would be 0.363 or

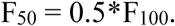

Since the significance of a change in muscle length or force will vary by context, we chose to present these data in two ways; as absolute changes normalized by maximum force capacity at 100% activation (F_100max_) and as relative changes between models. Absolute changes were plotted for all data points as a function of MTU compliance and passive length normalized to muscle length at 100% activation, L0_100_ (Fig. 3). Absolute changes relative to maximal force allowed us to quantify the contribution of each factor as well as painting a more detailed picture of the interaction between these factors and the conditions in which each is most influential. Yet, this approach can obscure the significance of changes at low activations. For instance, at 25% activation, a change in force capacity of 5% of maximum force between models may represent a relative change of anywhere from 20 to 80% of force capacity. To capture these relative effects, we again binned our data, but this time into much smaller bin sizes (steps of 5% of Lo, T/M ratio bins: 0.2 for low compliant muscles and 1 for high) and compared across models as described above (NCNS, CNS, CS). For each bin we calculated the number of samples in each bin and the percent of samples with a greater than 10% difference in force capacity between models. The 10% difference is an arbitrary cut-off below which we deemed the activation-compliance effect to be of less importance in dictating muscle function. The percent difference between muscle-tendon models was taken from the difference between the forces predicted by each model divided by the force predicted by the less complex model. For example, the percent difference between models with and without compliance (NCNS and CNS), was given by

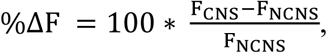

where F_CNS_ and F_NCNS_ are the forces predicted by the CNS model and NCNS models respectively. With this analysis we aimed to identify a threshold of MTU compliance or range of operating lengths where these factors can be safely ignored and the conditions in which simple models that exclude these factors lead to errors.

**Figure 3.**
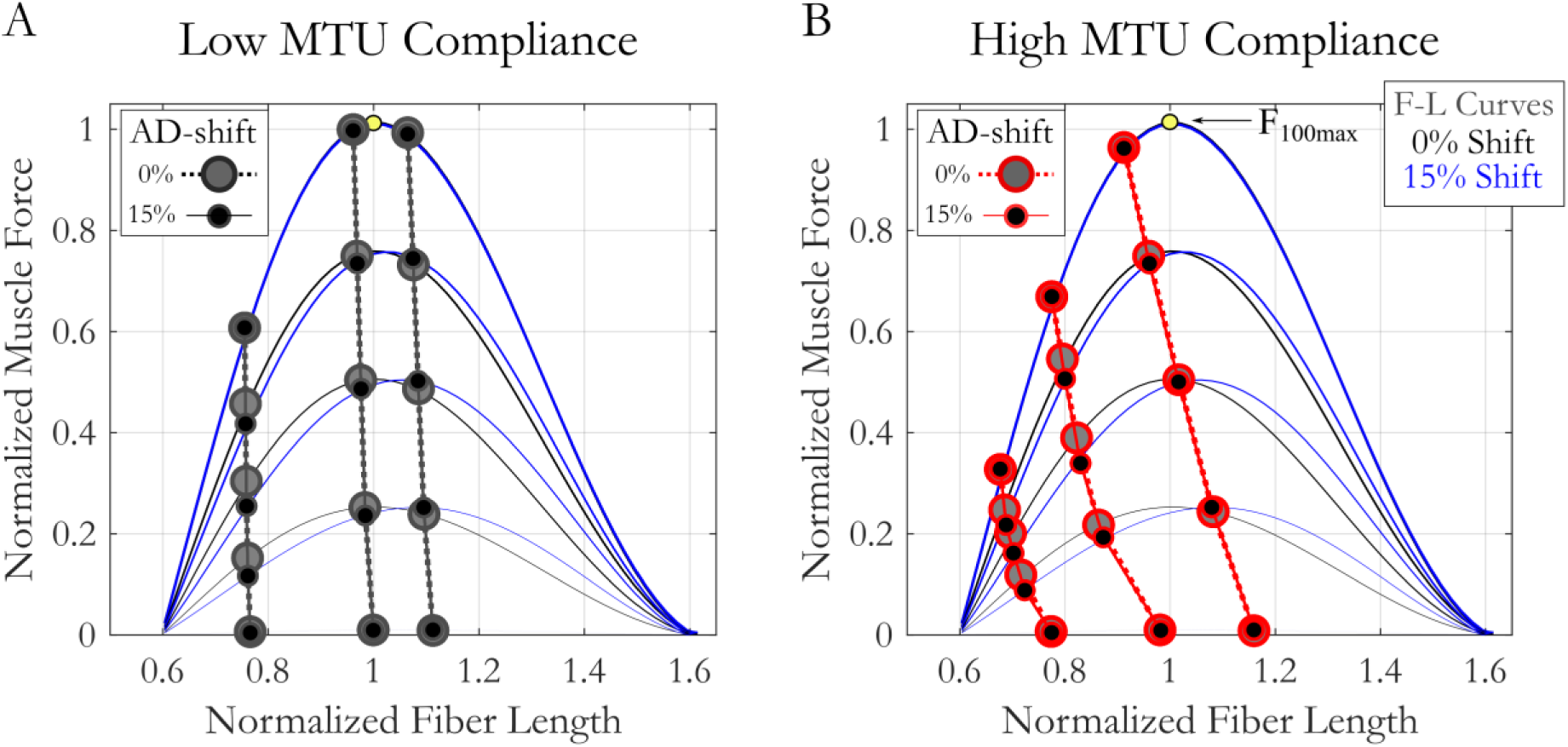
The influence of activation, MTU compliance, passive (pre-activation) muscle length and activation dependent shifts in L_0_ on average muscle length. Circles represent the mean value at the five different activation levels and circles from the same group are connected by lines across activation. L0_100_ is always a normalized fiber length of 1. Yellow circle marks the maximum force at 100% activation, F_100max._ On average, muscle length decreases with increasing activation and is significantly influenced by interactions between starting length, compliance and activation level. Muscles with low MTU compliance (A) change length with activation less than muscles with high compliance (B). While AD-shifts in L_0_ change operating length (difference between black and grey filled circles), they are not significant.

**3. What are the functional consequences of these factors?** Specifically, how do these activation dependent factors sum over many muscles acting across a joint to influence the posture for maximal force and moment capacity at low activation (25%) and how does this influence compare between a distal and proximal joint?

To explore these questions, we compared force and torque generated by the sum of muscles acting across the ankle and the hip. We chose to evaluate the muscles acting across the hip and ankle since on average they are composed of MTU’s with very different levels of compliance (T/M Ratio Hip:0.71, Ankle:7.82, Table 2) and did not overlap. We chose a consistent 25% activation to match the lowest level of activation used to evaluate our second question above and made it uniform to simplify the analysis. We simulated 25% activation of muscles acting across these joints for each model (NCNS, CNS, CS) through a sweep of hip and ankle postures with knee and TMP angle held constant at standing angles [Ankle and hip angle ranges as described in Fig. 1. Knee_flex_:-110 ^o^, TMP_flex_:-55 ^o^ (Henry et al., 2005)]. For each joint angle we extracted the total force and torque acting across each joint as well as the mean moment arm weighted by maximal muscle force, F_100max_ (Fig. 5). Maximum total muscle force and torque were found for each joint for each model as well as the corresponding joint angle (Table 4). For reference, we then calculated the percent difference between these joint angle-dependent values and the force or torque in the standing posture. The goal of this analysis was to provide an estimate of the combined influence of the activation-compliance dependent effects across many muscles at low activation.

## Results

**1. On average, how does activation level interact with MTU compliance, passive muscle length and/or activation dependent shifts in optimal fiber length to influence the operating range and force capacity of muscles?**

We found passive muscle length, activation and MTU compliance influenced how a muscle changes length with increasing activation, while activation dependent shifts in optimal fiber lengths did not (Fig. 3, Table 3). There were neither significant main nor interaction effects with activation-dependent shifts in L0. When models were re-run without including activation-dependent shifts in L0, we found 3^rd^ order interaction terms between passive muscle length, activation level and MTU compliance (Table 3). This implies that how muscle length changes with activation level differs between muscle of low and high compliance and that this relationship changes with the starting passive length of the muscle.

**Table 3.** Model parameters and results for linear mixed effects models with change in muscle length as dependent factor. Activation dependent shift in L0 (Shift) did not have significant main or interaction effect. Activation level, MTU compliance (TFR) and passive region of the force-length curve all interact to influence operating muscle length.

| Model | Shift*TMR*Limb*Act |  |  |  |  | TMR*Limb*Act |  |  |  |
| --- | --- | --- | --- | --- | --- | --- | --- | --- | --- |
|  | DF | Sum sq | Mean sq | F val | p-val | Sum sq | Mean sq | F val | p-val |
| Shift (1) | 1 | 0.02 | 0.02 | 17.4 | 0.15 |  |  |  |  |
| T/M Ratio (2) | 1 | 0.94 | 0.094 | 83.1 | 0.07 | 0.097 | 0.097 | 83.08 | 0.07 |
| **Limb (3)** | **2** | **13.07** | **6.53** | **5768.2** | **1.7e-4** | **6.8** | **3.3** | **2808.9** | **3.6e-4** |
| **Act (4)** | **4** | **83.87** | **20.96** | **18502.8** | **1.75e-8** | **41.9** | **10.5** | **8964.4** | **7.4e-8** |
| 1x2 | 1 | 0.017 | 0.167 | 14.70 | 0.01 |  |  |  |  |
| 1x3 | 2 | 0.016 | 0.008 | 7.03 | 0.01 |  |  |  |  |
| **2x3** | **2** | **5.95** | **2.97** | **2625.2** | **3.08e-4** | **3.04** | **1.5** | **1300** | **7.7e-4** |
| 1x4 | 4 | 0.018 | 0.005 | 4.07 | 0.1 |  |  |  |  |
| **2x4** | **4** | **39.83** | **9.96** | **8787.3** | **7.8e-8** | **19.98** | **4.99** | **4265.6** | **3.3e-7** |
| **3x4** | **8** | **14.49** | **1.81** | **1598.6** | **1.6e-10** | **7.5** | **0.94** | **804.5** | **2.6e-9** |
| 1x2x3 | 2 | 0.005 | 0.002 | 2.27 | 0.2 |  |  |  |  |
| 1x2x4 | 4 | 0.013 | 0.003 | 2.95 | 0.16 |  |  |  |  |
| 1x3x4 | 8 | 0.022 | 0.003 | 2.37 | 0.16 |  |  |  |  |
| **2x3x4** | **8** | **5.78** | **0.722** | **637.4** | **6.4e-9** | **3.02** | **0.38** | **322.1** | **9.5e-8** |
| 1x2x3x4 | 8 | 0.008 | 0.001 | 0.89 | 0.3 |  |  |  |  |

This complex interaction can be seen in Figure 3. While both high and low compliance muscles become shorter with increasing activation, high compliance muscles show a larger effect. The extent of the effect is also dependent on passive muscle length, with muscles with passive lengths on the descending limb of the force-length curve showing larger length changes with increasing activation for all levels of compliance (Fig. 3). This is explained on the basis that force capacity on the descending limb, and thus the compliance effect, increases as the muscle shortens towards L0.

**2. Under what combinations of these factors is the force generating capacity of an individual muscle meaningfully altered?** *How does compliance and activation dependent shifts in optimal fiber length alter muscle operating length and force capacity at high (100%) and low (25%) activation?*

### The influence of compliance

Increasing compliance increases how much a muscle shortens with increasing activation. At 100% activation (Fig. 4A), the change in length can be as high as 40% fiber strain while at 25% activation that decreases to 25% muscle fiber strain (Fig. 4B). At high activation, the starting passive length which results in the greatest length changes increases with MTU compliance (Fig. 4A, shifting from ~ 1.0 to 1.25 L0) while at low activations the passive length that produces the greatest length changes is less influenced by compliance (Fig. 4B, shifting only from ~ 1.0 to 1.05 L0). The consequence of these length changes on the force capacity depend on the passive muscle length, generally hampering force generation at short passive lengths and amplifying force capacity at long passive lengths (Fig. 4D). Though, it is interesting to note that change in force capacity does not linearly scale with passive muscle length. At the tails of the force-length curve (short and long extremes), the force effects are smaller due to the decreased force capacity. Thus, the passive length that results in the greatest change in muscle length does not occur at the extremes nor does it align with the length that results in the greatest change in force. The transition between decreasing and increasing force capacity as a result of activation-compliance effects occurs at longer passive muscle lengths for more compliant muscles. At 100% activation, the transition shifts from a normalized length of 1 at low compliance to 1.2 for high compliant muscles. The results at 25% activation, again, show similar patterns, but with less dramatic variations (Fig. 4E).

**Figure 4.**
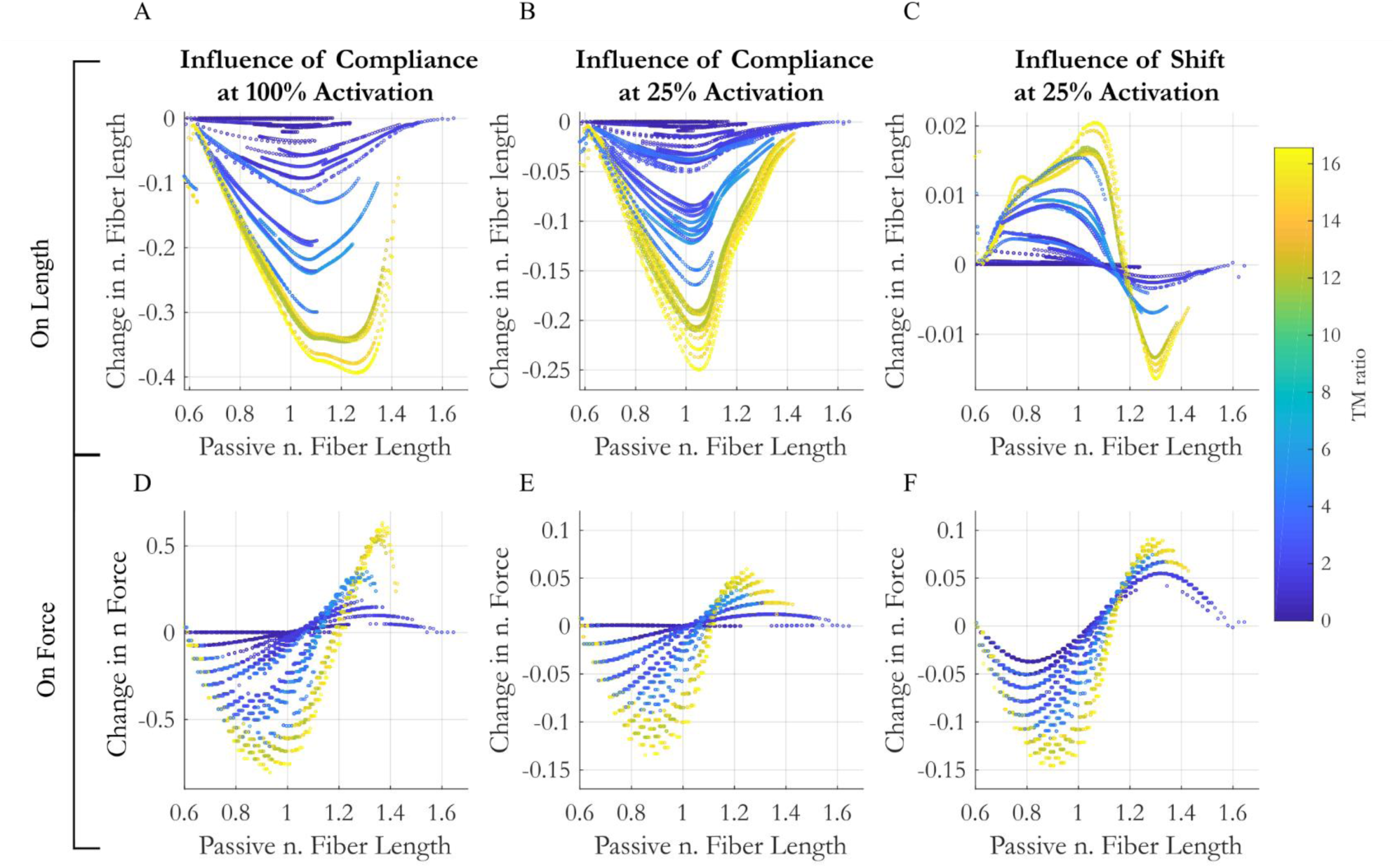
The influence of compliance (A,B,D,E) and an activation dependent shift (C&F) in the optimal fiber length on muscle length (A-C) and force (D-E) as a function of the passive (pre-activation) fiber length and MTU compliance (designated by color).

### The influence of activation dependent shifts in optimal fiber length

In comparison to the influence of compliance, activation dependent shifts in optimal fiber length result in much smaller changes in normalized muscle length. Muscle lengths change by at most 2% of L0_100_ (Fig. 4C). Unlike the influence of compliance, though, shifts in optimal fiber length result in longer as well as shorter muscles depending on passive muscle lengths. While intuition may suggest that activation-dependent shifts alone would not alter a muscle’s length, the variations are likely the result of muscles finding an equilibrium with the compliant tendon. For a muscle on the ascending limb of the force-length curve, for instance, a shift in the force-length curve to the right decreases its force capacity. This results in the muscle being able to stretch the tendon less. Thus, for any fixed muscle-tendon length, the equilibrium length of the muscle on the ascending limb is longer in the presence of activation-dependent shifts. The opposite effect can be seen on the descending limb (Fig. 4C), and again, these effects are amplified for more compliant MTU’s.

In contrast to the influence of compliance, which has little effect on the force generating capacity of low compliance muscles, activation-dependent shift in L0 influence the force capacity of muscles of all compliance levels, though the greatest effects are still seen in the most compliant MTU’s. Activation-dependent shifts in L0 can result in both decreased and increased force capacity, depending on passive muscle length and compliance. In general, muscles with passive lengths less than ~1.1 L0 decrease force capacity in the presence of activation-dependent shifts while muscles above a normalized length of 1.1 L0 increase force capacity. The magnitude of the change in force capacity with shift matches or exceeds the influence of compliance at 25% activation.

In summary, at high activation, compliance results in muscle strain up to 40%. These changes result in increases of force capacity for muscles on the descending limb of the force-length curve (up to 60% of maximum force, F_max_) and decreases on the ascending limb (of up to 80% of maximum muscle force capacity). Muscle force is most sensitive to compliance effects on the shallow ascending limb (Ls~0.7-0.9) or the middle descending limb (Ls~1.2-1.3) of the force-length curve. At low activations, though activation-dependent shifts in L0 result in only small changes in muscle length (at most 1-2% strain), they have a larger effect on force capacity than compliance (Fig. 4F vs Fig. 4E) since the peak of the force-length curve is changing relative to the muscle length. While the influence of activation-dependent shifts in L0 on force capacity decreases with compliance, for muscles with no tendon, they can be as high as 5% of maximum force capacity.

### Which combinations of these factors alter the relative force capacity by more than 10%?

#### Influence of compliance

The range of passive lengths that result in a change of force capacity increases with compliance and activation, as expected. For the most compliant MTU’s, 88% of starting passive muscle lengths result in a change in force capacity that is greater than 10% (Fig. 5A). While our data does not allow us to fully quantify the space, it does allow us to isolate regions of high and low sensitivity to compliance. For instance, at both 100 and 25% activation, *all* MTU’s with a T/M ratio of as low at 0.8 show a meaningful (>10%) influence of compliance on force capacity when the muscle’s initial passive length is between 0.7 and 0.75 L0 (Figs. 5A&B). Thus, even low compliance MTU’s have a region of operating lengths where compliance meaningfully influences force capacity. There also exists a trough of influence where compliance and AD shifts of L0 have little influence on force capacity. The location of this trough varies with MTU compliance, though, shifting slightly to longer muscle lengths with increasing compliance. For example, MTU’s with a T/M ratio of 1.2 show no change in force capacity with compliance when operating at 1.05 L0, while the passive length that has the smallest influences on the force capacity of the most compliant muscles is closer to 1.15 at 25% activation and 1.25 at 100% (Fig. 5C).

**Figure 5.**
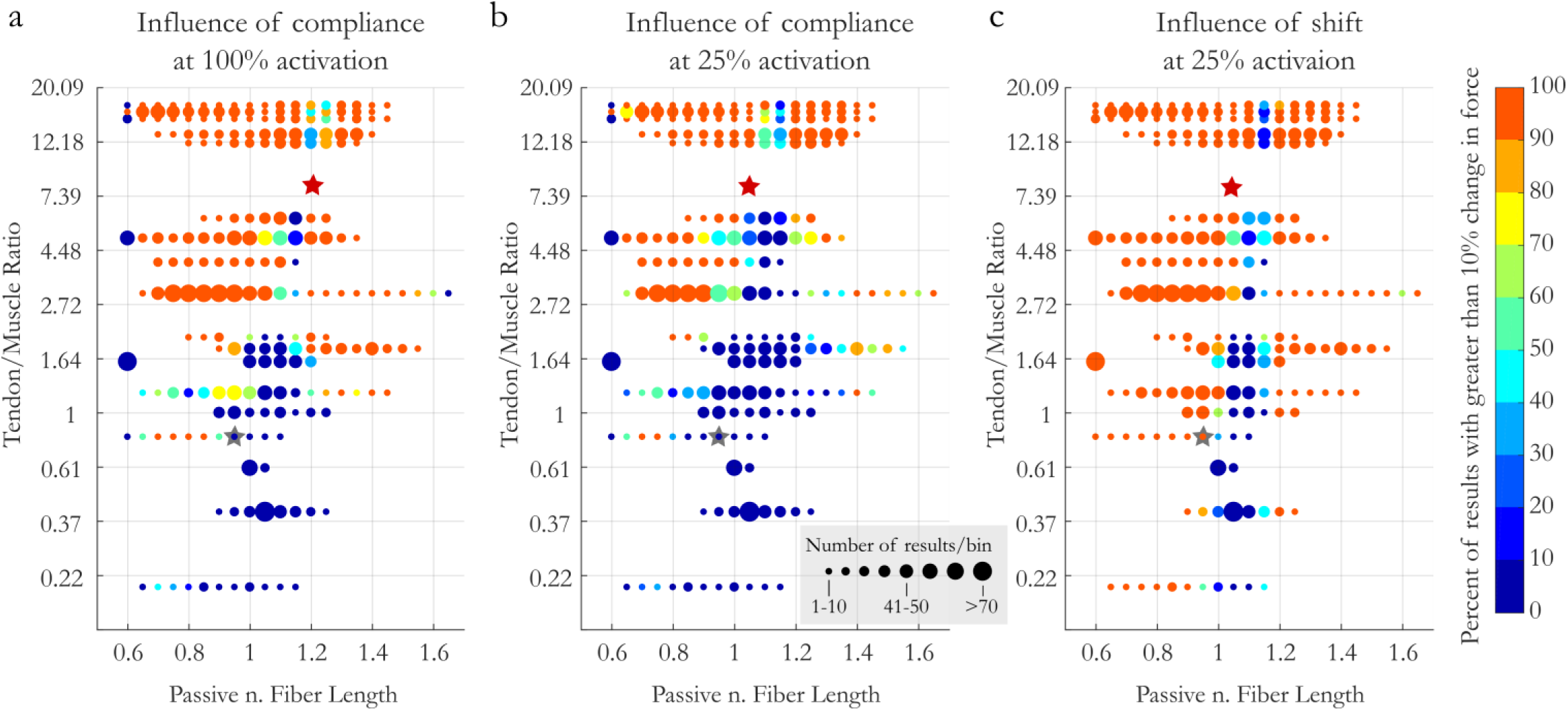
The influence of compliance (at 100% (a) and 25% activation (b)) and AD shifts in L0 on force production [at 25% activation,(c)] as a function of MTU compliance and passive (pre-activation) muscle length. Color designates the percent of results that saw a greater than 10% change in force production between models. Circle size represents the number of samples per bin. Y-axis is non-linear to ease visualization of results at low MTU compliance. Stars represent average compliance and operating length of the muscles acting at the hip (grey) and ankle (red) in CNS (b) and CS (c) conditions

#### Influence of activation-dependent shifts in L0

Unlike the influence of compliance, activation-dependent shifts alter the force capacity of muscles of all levels of compliance. While compliance had very little influence on the force capacity for muscles with very little tendon, activation-dependent shifts in L0 resulted in changes in force capacity of over 10% for muscles with no tendon (T/M ratio = 0) across half of the operating range, primarily at lengths on the ascending limb. Again, the influence increased with increasing compliance, such that for the most compliant muscles activation-dependent shifts in L0 resulted in changes in force capacity for all muscles across 95% of the possible initial passive lengths of the muscle. The trough of influence for activation-dependent shifts in L0 followed similar patterns to that for compliance, with minimal effect on force for stiff MTU’s at lengths of 1.05 and for compliant muscles at 1.15.

To summarize, at high activation compliance considerably changes the force capacity for muscles with a T/M ratio over 2 at nearly all passive lengths (95%). While change in force capacity increases with compliance, all muscles with low compliance (i.e. T/M ratio as low as 0.7) are also affected when passive muscle lengths are in the middle of the ascending limb (Lp: 0.7-0.85). At 25% activation the results are similar but with decreased magnitude. Activation-dependent shifts considerably influence the force capacity of all muscles on the ascending limb of the force-length curve. For all of these effects, there is a trough of influence near or slightly above L0 where activation does not alter force capacity appreciably.

**3. How do these compliance and activation dependent factors influence the posture which maximizes the force generating or moment capacity?**

#### Influence of compliance

As expect, compliance has less of an effect on the force capacity for muscles acting at the hip than for more distal muscles acting at the ankle. At the hip, the postures that maximizes force and moment capacity change little with compliance (4° and 6°, respectively). In contrast, at the ankle the postures that maximizes force and moment capacity change by 26° and 20°, respectively (Figs. 6a&b, Table 4). Likewise, we see very little difference in force capacity at the hip between models that do and do not include compliance in a standing posture (<-6%, blue vs. grey solid lines at 50° Fig. 6a). When muscle moment arms are taken into account we observe a reduction in moment capacity (−15%) compared to maximum values at a more extended posture. At the ankle, muscles operating on the descending limb of the force-length curve in a standing posture benefit from the change in length with compliance such that they approach L0_100_ and increase their force capacity by 27% and joint moment by +23% (CNS model; blue vs grey dashed lines at 98°, Fig. 6a).

**Figure 6.**
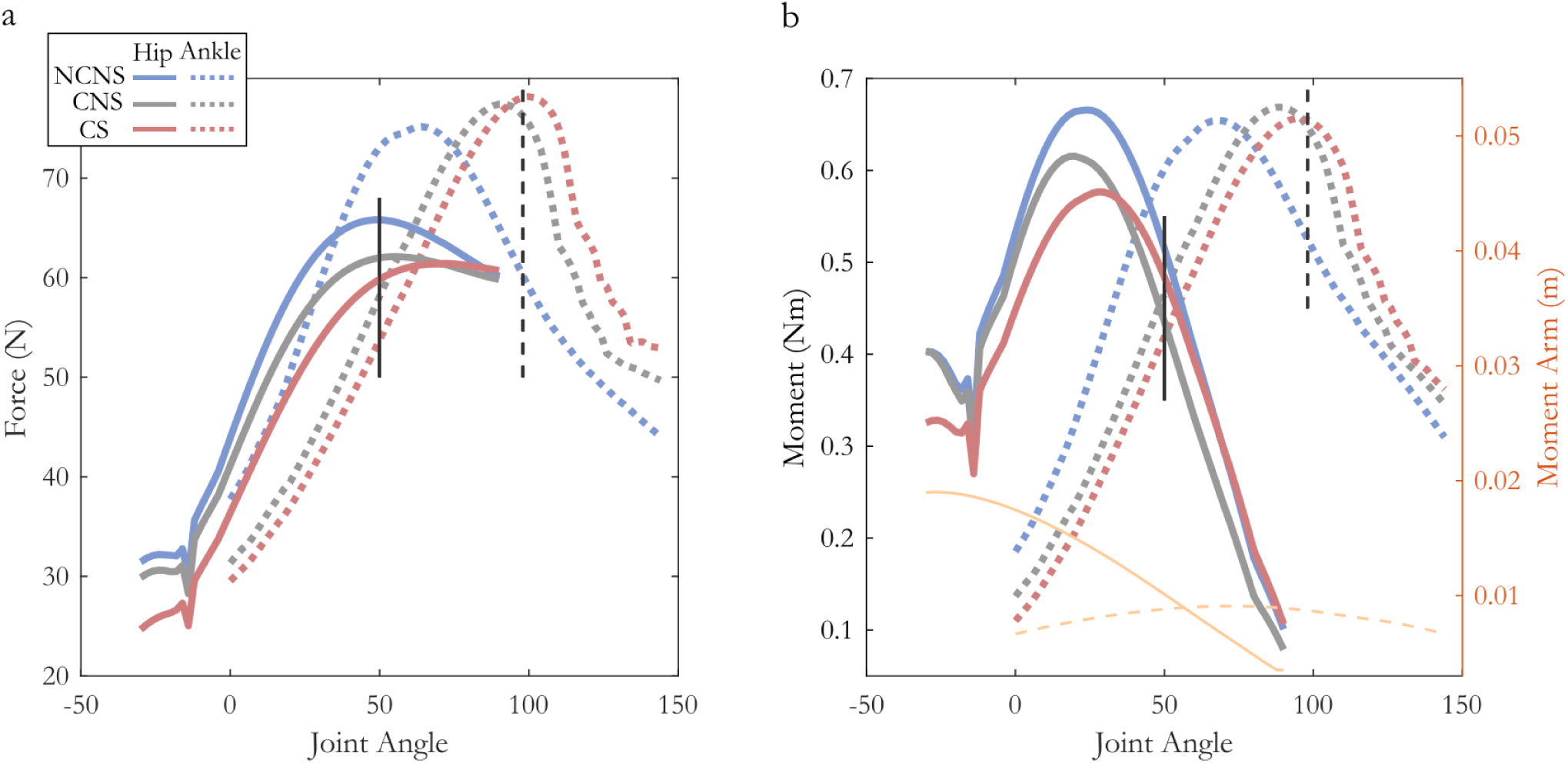
Joint angle vs total force (a) and moment (b) acting across the hip (solid lines) and ankle (dashed lines) under three different muscle-tendon models. Average weighted moment arms for the hip (solid) and ankle (dashed) depicted in orange. While at the hip, the posture that produces maximum force and moment does not vary much between models (<9 ^o^), the maximum force and moment capacity decreases with both compliance and Ad-shift (~13%). In contrast, at the ankle the magnitude of maximum force and torque varies little between models (<3%), but the posture at which this maximum occurs varies by almost 40°.

**Table 4.** A comparison of functional influence of three different muscle-tendon models: No compliance No Shift (NCNS), Compliance but No Shift (CNS) and Compliance and Shift (CS). For each model, the angle at maximum force (Fmax^o^), average normalized fiber length (nFL), and total force (F) and moment (M) across the joint are listed for both the hip (low compliance) and ankle joints (high compliance). Compliance and shifts in L0 decrease the force and torque capacity at the hip while increasing them at the ankle.

|  | Hip T/M ratio: 0.71 |  |  |  |  |  |  | Ankle T/M ratio 8.46 |  |  |  |  |  |  |
| --- | --- | --- | --- | --- | --- | --- | --- | --- | --- | --- | --- | --- | --- | --- |
| | Max Values | | | | Standing $50^\circ$ | | | Max Values | | | | Standing $98^\circ$ | | |
|  | Force |  | Moment |  | nFL | F(N) | M (Nm) | Force |  | Moment |  | nFL | F(N) | M (Nm) |
| | Mag | $^\circ$ | Mag | $^\circ$ | | | | Mag | $^\circ$ | Mag | $^\circ$ | | | |
| NCNS | 65.8 | 50 | 0.67 | 24 | 0.97 | 65.8 | .52 | 75.2 | 64 | 0.65 | 68 | 1.21 | 60.2 | .53 |
| CNS | 62.1 | 56 | 0.62 | 20 | 0.95 | 62.0 | .44 | 77.4 | 90 | 0.67 | 88 | 1.06 | 76.3 | .65 |
| CS | 61.4 | 72 | 0.58 | 28 | 0.95 | 59.8 | .48 | 78.2 | 100 | 0.66 | 96 | 1.05 | 78.1 | .60 |

#### Influence of activation-dependent shifts in L0

The influence of activation-dependent shifts shows a different pattern than that of compliance. Both the hip and ankle are equally influenced by activation dependent shifts in L0. At the hip, the postures that maximizes force capacity and moment change by 16° and 8° when activation-dependent shifts in L0 are included in the model, while the ankle the postures that maximizes force capacity and moment change by 8° and 10° (Figs. 6a&b respectively, Table 4).

In a standing posture, muscles acting act the hip, since they operate on the ascending limb, are shifted further away from optimal length when an activation-dependent shift in L0 is added to the model and show a decrease in force and moment capacity (Δforce:-4%, Δmoment:-9%, Fig. 6A solid red line). The ankle, though, operates on the plateau of the force-length curve (~1.06) and its force capacity increases slightly (Δforce:1.5%, Δmoment:+9%) when an activation-dependent shift in L0 is added to the model (Fig. 6A, dashed red line).

In summary, at the hip, compliance decreases force capacity slightly more than AD shifts in L0 (−5% and −4%, respectively) in a standing posture, with both effects being small. At the ankle, compliance increases the force generated at the ankle substantially (+26%), while activation-dependent shifts have a much smaller effect (+2%). Similar effects are seen for moment generating capacity.

## Discussion

The force-length relationship is one of the key components affecting a muscle’s capacity to produce force. And while there are multiple factors that are known to influence a muscle’s force-length operating range, their contribution and possible interactions have not been well quantified. Here we tested the effect of, and interaction between, muscle tendon compliance, activation and activation-dependent shifts in L0 on muscle operating length and force during simulated fixed-end contractions in architecturally diverse muscles across a multitude of conditions.

### General effect of compliance and activation-dependent shifts in L0 on muscle length and force capacity

One of the overarching goals for this study was to identify general principles (if any) that describe how compliance and activation-dependent shifts in L0 influence muscle operating lengths. While the relationships were quite complex (see following section), we reveal several notable patterns. First, we found that in this species, on average, most muscles operate on the plateau or ascending limb of the force-length curve irrespective of the level of tendon compliance or activation-dependent shifts in L0. These averages incorporate data across the total possible *in vivo* length range of the muscles, including both the full range of passive lengths and activation levels. Because our analysis spans muscle force levels from near zero to maximum values, we do not expect that force-length ranges would deviate from this in dynamic contractions. The exception is possibly eccentric contractions under maximal activations where force can exceed F_max_ (Herzog et al., 2006). This comprehensive simulation of muscle operating lengths corroborates much of the literature showing that muscles typically do not operate on the descending limb of the force-length curve (Burkholder et al., 2001; Holt and Azizi, 2016; Rubenson et al., 2012). This provides further support for the idea that an animal’s neuro-musculoskeletal structure is most often organized to operate on the ascending limb or plateau regions where muscles are inherently more stable and less susceptible to stretch-induced injury (Herzog et al., 1992, 1991; Lindstedt et al., 2001; Lutz and Rome, 1996; Morgan et al., 2002).

Our results also show that the most common effect of compliance is a leftward shift on the force-length curve leading to shorter muscle lengths and a decrease in force capacity. Since muscles, on average, operate on the ascending limb, increasing activation shortens muscles further below L0 as the tendon stretches with greater force generation. The exception is for muscles that start at passive lengths longer than L0 Under this condition, the result is an increase in force capacity as the compliant muscle’s fibers shorten towards L0, though some muscle lengths can shorten beyond L0, first increasing and then decreasing force capacity. It is also notable that the average effect of activation-dependent shifts in L0 generally mirror those described above for compliance, although with much smaller effects. Thus, on average, the effects of compliance dominates activation-dependent changes in operating lengths.

Finally, by comparing results that average across muscles acting at the hip (low compliance) with those acting at the ankle (high compliance), we show that more compliant tendons will generally amplify the effects described above. Given the proximal-distal gradient of compliance, this analysis suggests that proximal muscles will show little activation-compliance effect whereas distal muscles will be more significantly influenced. In summary, if specific knowledge of a muscle’s fiber, tendon and joint characteristics are lacking, the average data provide a cursory prediction of the most likely influence of compliance and activation-dependent shifts in L0 on muscle force-length operating ranges. As muscles increase activation, they will shorten further away from L0 and generate less force than would be predicted by a linear force-activation relationship. This effect increases with compliance and muscle passive length (up to L0). Activation dependent shifts in L0 amplify this effect, but only minimally. However, these generalizations should be used with caution because, as we will detail in the following sections, they are error prone and can obscure the highly variable effect of compliance and/or activation-dependent shifts in L0 on functionally-relevant changes in length and force capacity.

### Under what conditions are the compliance and activation-dependent shifts in L0 meaningful?

While the analysis of average behavior paints results with a wide brush, it fails to capture the complex interactions between the muscle’s starting passive length, compliance and activation-dependent shifts in L0. While there are trends in these interactions, it is important to emphasize that they are not broad. These relationships are very sensitive to a muscle’s passive length and have large continuous gradients that shift with MTU compliance. Thus, small changes in the initial passive length can result in a large change in activation-compliance-dependent force capacity that generalizations fail to capture. For instance, while averages suggest that activation-dependent shifts in L0 have little influence on force capacity, this more detailed analysis shows that they can decrease or increase force capacity by as much as 15% of F_100max_, resulting in a relative change of over 60% in force capacity at low activations. Likewise the average analysis underrepresented the extent of compliance on muscle length, which can be drastic, resulting in changes of up to 40% of L0. This is commensurate with the largest muscle strains seen in dynamic, high power movements (Askew and Marsh, 2002). Here we show this in a fixed-end ‘isometric’ contraction. Further, while the ‘average’ analysis concluded only a small effect of activation on muscle length in non-compliant muscles, this more detailed analysis highlighted that there is no abrupt threshold of MTU compliance below which the effects disappear. Rather there is a non-linear continuous gradient with compliance that varies across muscle operating lengths. There are regions of the force-length curve in which muscles are most sensitive to compliance and there are troughs of influence where these factors have little effect on force capacity. These regions are hard to predict, though, because the magnitude and location of these areas of high and low sensitivity shift with compliance level. Thus, rather than being able to provide an accurate rule of thumb for how and when compliance will alter a muscle’s force generating capacity, we identify conditions in which the effect is most drastic, the variables that are most influential, and provide a guide for the possible errors in predicting muscle function if muscle-tendon properties are not known.

While, on average, changes in muscle length were greatest for muscles starting at passive lengths closer to optimal fiber length, L0, the influence of compliance on force capacity does not follow suit. The length at which we see the greatest change in force shifts with MTU compliance. On the ascending limb, stiff MTU’s show the greatest drop in force at the shortest passive lengths (dark blue regions of Fig. 4d), while compliant muscles see the greatest change when muscles start at passive lengths just short of optimal fiber lengths (yellow regions of Fig. 4d). While it is fair to say that compliance will always decrease the force generating capacity of muscles with passive lengths on the ascending limb, the non-linear interactions between activation and compliance make any approximations of the magnitude of the effect potentially inaccurate. Depending on passive muscle length, the influence of compliance could decrease a muscle’s force capacity on the ascending limb anywhere from zero to 80%. While these numbers are striking, this isn’t even the region of the force-length curve that is most sensitive to the effects of compliance. The region of most variability extends from just short of the plateau region of the force-length curve and down a portion of the descending limb. For muscles within this region, at 100% activation compliance can result in either a decrease of or increase in force capacity of well over 50% of F_100max_, depending on the passive length. It is important to emphasize that these conditions in which we see the greatest influence of activation on operating length are not unusual. Within this region of highest variability are conditions we typically regard as ‘optimal’, on or near the plateau. Thus, surprisingly, across the region of the force-length curve we typically view force as relatively insensitive to length changes (i.e. the plateau), we find the area of greatest sensitivity to activation-compliance effects. In summary, compliance results in activation dependent changes in the operating length and force capacity of muscles, the maximum values of which increase with increasing compliance. On the ascending limb, force capacity decreases with increasing compliance and passive muscle length. On the plateau or descending limb, the story is much more complex and force capacity can either increase or decrease depending on compliance and initial passive muscle length. Thus, at a muscle function level we show that even in “isometric” conditions, the muscle-tendon unit is really ‘dynamic’ with changes in muscle length, force capacity and activation requirements. One cannot accurately assume force increases proportionally with activation; the activation-force relationship is far from linear. Instead, it is likely a complex interaction of activation, muscle passive length and MTU compliance. This interaction, although not well understood, suggests great computational complexity for neural control of muscle force production.

The influence of activation-dependent shifts in L0 on force capacity generally follows a similar pattern as described for compliance, increasing with increasing compliance. Again, this can be explained by the dynamic nature of these interactions. The changes in force capacity that come with a shift in L0 result in larger length changes in more compliant MTU’s, which result in further force changes. While these effects can be small in magnitude (less than 5% of maximum force capacity for T/M ratio < 2), at low activation, this can represent a relatively large change in force capacity (over 20%). In fact, at low activation, the influence of activation-dependent shifts in L0 is as important, if not more important, than compliance effects in determining a muscle’s force capacity. Further, there is only a narrow region of muscle lengths that result in ‘no influence’; thus, the contribution of activation-dependent shifts in L0 are more ubiquitous than our average analysis suggested, substantially altering the force capacity across a wide range of MTU compliance and operating lengths at low activations. In summary, this analysis allows us to provide information on what the most important variables are for accurately predicting muscle force. Compliance is more important than activation-dependent shifts in L0 at high activation, while both significantly alter the force capacity at low activation levels.

This has implications for musculoskeletal modeling and can provide a guide for what the possible errors are in predicting muscle function if muscle-tendon properties are not known. Models that assume non-compliant tendons could be over or underestimating muscle force drastically. Specifically, this simplification would most often result in over estimation of force generating capacity with muscles operating on the ascending limb and under estimation on the descending limb. This is true, even within the less compliance range of human MTU’s [lower limb av. 3.06 (Arnold and Delp, 2011)]. At a T/M ratio of 3, for instance, our analysis shows that for all but a narrow band of passive lengths, muscles show significant force deficits with compliance. More striking is that even muscles with relatively short tendons (tendons shorter than the muscle optimal fiber length, T/M rations lower than one) have regions of passive lengths on the ascending limb where compliance result in significant changes in force capacity. This implies that even low levels of compliance can significantly alter force capacity in some conditions and that models that ignore or do not accurately define compliance should be interpreted cautiously. More optimistically, muscle models that include compliance but ignore activation-dependent shifts in L0 should capture the most dominant factors when simulating muscles at high activations. But in cases where muscles are at submaximal activation (i.e. many activities), ignoring activation-dependent shifts in L0 could lead to errors that mirror those of non-compliant models; in general forces on the ascending limb are overestimated and those on the descending limb underestimated.

### What are the functional consequences of compliance / activation-dependent shifts in L0?

While the major focus of this paper is to understand in detail the influence of compliance and activation-dependent shifts in L0 on muscle operating lengths and force capacity, we also make a first pass at assessing their possible functional influence. We do this in a task where our ‘static’ analysis is most applicable, namely in a standing posture.

First, consistent with results from our other analyses, we find that force in the hip muscles, which shorten down the ascending limb, decrease when both compliance and activation-dependent shifts in L0 are included in the muscle models, whereas forces in the ankle muscles, which shorten up the descending limb, increase with the addition of activation dependent effects. This is also consistent with our finding that activation dependent effects generally have a larger influence on more compliant MTU’s. Models that include both activation dependent effects have a greater influence of ‘optimal force-generating posture’ at the ankle than at the hip. Surprisingly, though, the influence of activation-dependent shifts in L0 are greater for muscles acting at the hip than those acting at the ankle, despite our results which show that activation-dependent shifts generally have larger effects for more compliant muscles. This emphasizes two take home points from this study. 1) Muscles with very little tendon are not immune to non-linear activation dependent effects, and 2) as stated previously, the magnitude of activation-dependent effects are very sensitive to the lengths at which the muscles operate. Here, the muscles acting across the ankle fall into a trough of influence in the plateau region where activation-dependent shifts in L0 have little impact (red star, Fig. 5c), while the muscles acting at the hip operate on the ascending limb in a region of relatively high influence (grey star Fig. 5c). Thus, again, we emphasize how highly sensitive activation-dependent effects are to muscle passive lengths and how broad generalizations can lead to erroneous conclusions.

In a previous study, we also asked whether animals utilize postures that maximize torque capacity during walking and/or running. These analyses concluded that they did not, but were based on simpler muscle models that did not incorporate activation dependent effects [compliance or activation-dependent shifts in L0 (Hutchinson et al., 2015)]. This previous study also did not separate the roles of length-dependent muscle force capacity and muscle moment arms in dictating the effect of joint posture on moment capacity. Several experiments have shown that peak isometric muscle force and moment arms do not necessarily coincide with the joint angle that generates peak muscle or joint moments (Hoy et al., 1990; Lieber and Boakes, 1988a). Whether muscle force or moment arms have the largest contribution to peak moment is variable. For example, for the same muscle (bi-articular frog semitendinosus), force capacity dictates the muscle’s peak knee joint moment (Lieber and Boakes, 1988b), but the muscle’s moment arm dictates the moment-angle profile at the hip (Lieber and Shoemaker, 1992). How length-dependent force capacity and muscle moment arms influence observed postures remains poorly understood.

If we assume that models that incorporate both compliance and activation-dependent shifts in L0 accurately capture muscle dynamics in this species and a conservative 25% muscle activation, we do find some evidence for muscle-tendon mechanics and posture operating in concert. Notably, we find that at the ankle, animals stand with a posture that maximizes both force and joint moment, accommodating the strain in the muscle that occurs due to the high compliance of its MTUs. These could offer support to the long-held theory that muscles operate at a length that minimize force losses arising from their force-length relationship (Azizi, 2014; Azizi and Roberts, 2010; Lieber and Ward, 2011; Rubenson et al., 2012; Smith et al., 2007).

However, the story is complex. At the hip, we see a difference of ~45° between postures that maximize torque and force, with the animal choosing a posture halfway between. Standing posture is ~22° more extended than optimal for force production and ~22° more flexed than the posture that maximizes joint torque. This symmetry obscures the functional significance of these differences, though. Since force capacity is less sensitive to changes in joint angle, a postural difference of 22° results in a loss of only 20% in force capacity but decreases joint moment by nearly 40%. It is interesting to note that adopting a posture that would increase moment capacity at the hip (more erect) would also decrease the moment necessary to oppose gravity. Thus, a less flexed hip angle could be more economically supported, yet is not adopted. While 25% activation may overestimate muscle activations during standing (Rubenson and Marsh, 2009), a lower activation level will only amplify these effects. Thus, this study, along with our previous work and those of others, hint that there are likely many factors that influence preferred posture and torque capacity is not universally prioritized. While our results are among the first to systematically link length-dependent force capacity with posture, whether this specific mechanical constraint strongly dictates posture is still far from clear.

## Limitations and Future directions

While we have done these analyses assuming a particular level of activation dependent shift of the force-length curve, several studies have suggested that the level of shift my vary by species and may range from having no influence (de Brito Fontana and Herzog, 2016), to as large as 60% shift (Holt and Azizi, 2014). Our analysis of a 15% shift level was not intended to produce a definitive quantification of its influence, but as a first pass at evaluating the possible contribution and interaction effects. These analyses could be repeated for different levels in the AD shift in L0 and in dynamic conditions that include force-velocity effects to quantify how this influences the general conclusions. Likewise, care should be taken in extrapolating our results for species with different tendon properties for a similar muscle/tendon ratio could result in very different force and length changes. Regardless of these limitations, this analysis highlights the advantages of exploring these questions via musculoskeletal modeling, an approach which allows the generation of thousands of data points to elucidate patterns and trends that are not visible from studies of one or two muscles alone.

## Supporting information

Supplemental Methods Model Development

## Author Contributions

S.M.C. and J.R. contributed to the conception and design of the study. S.M.C., K.E., S.L.D. and J.R. were the primary authors involved in the development of the musculoskeletal model and S.M.C developed the muscle simulations. M.C., R.L.M. and J.R. were the primary authors involved in the collection and analysis of experimental data. All authors contributed critically to data interpretation. S.M.C and J.R. drafted the initial manuscript and S.M.C., K.E., R.L.M, S.L.D. and J.R. contributed to editing the manuscript. S.M.C and J.R. contributed to figure preparation.

## Acknowledgements

The authors would like to thank Andrew Wong for his assistance with muscle dissection and muscle-tendon path digitizing and spatial transformations and Hardik Sanghvi for his assistance with model implementation. This study benefited from helpful discussions with the Stanford Neuromuscular Biomechanics Lab fellows and students, in particular Frank ‘Clay’ Anderson, Thor Besier, and Ajay Seth. Robert Siston helped develop the optical tracking software and interface used to digitize muscle-tendon paths.

## Data Availability

Upon acceptance, data will be made available on Data Dryad and model will be made public on SimTK.

## References

Arnold, EM, Delp, SL. 2011. Fibre operating lengths of human lower limb muscles during walking. Philosophical Transactions of the Royal Society B: Biological Sciences, 366(1570): 1530–1539.

Arnold, EM, Hamner, SR, Seth, A, Millard, M, Delp, SL. 2013. How muscle fiber lengths and velocities affect muscle force generation as humans walk and run at different speeds. Journal of Experimental Biology, 216(11): 2150–2160.

Askew, G, Marsh, R. 2002. Muscle designed for maximum short-term power output: quail flight muscle. Journal of Experimental Biology, 205: 2153–2160.

Azizi, E. 2014. Locomotor function shapes the passive mechanical properties and operating lengths of muscle. Proceedings. Biological Sciences / The Royal Society, 281(1783): 20132914.

Azizi, E, Roberts, TJ. 2010. Muscle performance during frog jumping: influence of elasticity on muscle operating lengths. Proceedings of the Royal Society B: Biological Sciences, 277(1687): 1523–1530.

Bates, D, Maechler, M, Bolker, B, Walker, S. 2015. Fitting Linear Mixed-Effects Models Using {lme4}. Journal of Statistical Software, 67(1): 1–48.

Blix, M. 1894. Die Lange und die Spannung des Muskels. Skandinavisches Archiv Fur Physiologie, 5(1): 173–206.

Buchanan, TS, Lloyd, DG, Manal, K, Besier, TF. 2004. Neuromusculoskeletal modeling: Estimation of muscle forces and joint moments and movements from measurements of neural command. Journal of Applied Biomechanics, 20(4): 367–395.

Burkholder, TJ, Lieber, RL, Burkholder, TJ. 2001. Sarcomere length operating range of vertebrate muscles during movement. Journal of Experimental Biology, 204(Pt 9): 1529–1536.

Carr, JA, Ellerby, DJ, Marsh, RL. 2011. Differential segmental strain during active lengthening in a large biarticular thigh muscle during running. Journal of Experimental Biology, 214(Pt 20): 3386–95.

Cromie, MJ, Sanchez, GN, Schnitzer, MJ, Delp, SL. 2013. Sarcomere lengths in human extensor carpi radialis brevis measured by microendoscopy. Muscle and Nerve, 48(2): 286–292.

Daley, MA, Biewener, AA. 2011. Leg muscles that mediate stability : mechanics and control of two distal extensor muscles during obstacle negotiation in the guinea fowl. Philosophical Transactions of the Royal Society B: Biological Sciences, 366: 1580–1591.

Daley, MA, Usherwood, JR, Felix, G, Biewener, AA. 2006. Running over rough terrain : guinea fowl maintain dynamic stability despite a large unexpected change in substrate height. Journal of Experimental Biology, 209: 171–187.

de Brito Fontana, H, Herzog, W. 2016. Vastus lateralis maximum force-generating potential occurs at optimal fascicle length regardless of activation level. European Journal of Applied Physiology, 116(6): 1267–1277.

Delp, SL, Loan, JP. 1995. A graphics-based software system to develop and analyze models of musculoskeletal structures. Computational Biological and Medicine, 25(1): 21–34.

Ellerby, DJ, Henry, HT, Carr, JA, Buchanan, CI, Marsh, RL. 2005. Blood flow in guinea fowl Numida meleagris as an indicator of energy expenditure by individual muscles during walking and running. Journal of Physiology, 2: 631–648.

Ellerby, DJ, Marsh, RL. 2006. The energetic costs of trunk and distal-limb loading during walking and running in guinea fowl Numida meleagris II. Muscle energy use as indicated by blood flow. Journal of Experimental Biology, 209: 2064–2075.

Ellerby, DJ, Marsh, RL. 2010. The mechanical function of linked muscles in the guinea fowl hind limb. Journal of Experimental Biology, 213(13): 2201–2208.

Fukunaga, T, Ichinose, Y, Ito, M, Kawakami, Y, Fukashiro, S. 1997. Determination of fascile length and pennation in a contracting human muscle in vivo. Journal of Applied Physiology, 82(1): 354–8.

Gordon, AM, Huxley, AF, Julian, FJ. 1966. The variation in isometric tension with sarcomere length in vertebrate muscle fibres. Journal of Physiology, 184(1): 170–192.

Gordon, JC, Rankin, JW, Daley, MA. 2015. How do treadmill speed and terrain visibility influence neuromuscular control of guinea fowl locomotion? Journal of Experimental Biology, 218: 3010–3022.

Hawkins, D, Bey, M. 1997. Muscle and tendon force-length properties and their interactions in vitro. Journal of Biomechanics, 30(1): 63–70.

Henry, HT, Ellerby, DJ, Marsh, RL. 2005. Performance of guinea fowl Numida meleagris during jumping requires storage and release of elastic energy. Journal of Experimental Biology, 208: 3293–3302.

Herzog, W, Lee, EJ, Rassier, DE, Schappacher, G, DuVall, M, Leonard, TR, Herzog, JA. 2006. Residual Force Enhancement Following Eccentric Contractions: A New Mechanism Involving Titin. Physiology, 31(3): 300–312.

Herzog, W, Leonard, TR, Renaud, JM, Wallace, J, Chaki, G, Bornemisza, S. 1992. Force-length properties and functional demands of cat gastrocnemius, soleus and plantaris muscles. Journal of Biomechanics, 25(11): 1329–1335.

Herzog, W, Read, LJ, ter Keurs, HEDJ. 1991. Experimental determination of force-length relations of intact human gastrocnemius muscles. Clinical Biomechanics, 6(4): 230–238.

Holt, NC, Azizi, E. 2014. What drives activation-dependent shifts in the force-length curve? Biology Letters, 10(9): 20140651-.

Holt, NC, Azizi, E. 2016. The effect of activation level on muscle function during locomotion: are optimal lengths and velocities always used? Proceedings of the Royal Society B: Biological Sciences, 283(1823): 20152832.

Hoy, M, Zajac, E, Gordon, E. 1990. A musculoskeletal model of the human lower extremity: The effect of muscle, tendon, and moment arm on the moment-angle relationship of musculotendon actuators at the hip, knee, and ankle. Journal of Biomechanics, 23(2): 157–169.

Hutchinson, JR, Rankin, JW, Rubenson, J, Rosenbluth, KH, Siston, RA, Delp, SL. 2015. Musculoskeletal modelling of an ostrich (Struthio camelus) pelvic limb: influence of limb orientation on muscular capacity during locomotion. PeerJ, 3: e1001.

Huxley, AF, Simmons, RM. 1971. Proposed mechanism of force generation in striated muscle. Nature, 233: 533–8.

Lakatta, EG, Jewell, BR. 1977. Length-dependent activation: its effect on the length-tension relation in cat ventricular muscle. Circulation Research, 40(3): 251–7.

Lemos, RR, Epstein, M, Herzog, W. 2008. Modeling of skeletal muscle: The influence of tendon and aponeuroses compliance on the force-length relationship. Medical and Biological Engineering and Computing, 46(1): 23–32.

Lieber, R, Brown, CG. 1992. Sarcomere length Joint angle relationships of seven frogs hindlimb muscles. Acta Anatomica, 145: 289–295.

Lieber, RL, Boakes, JL. 1988a. Muscle force and moment arm contributions to torque production in frog hindlimb. American Journal of Physiology, 254(6 Pt 1): C769–72.

Lieber, RL, Boakes, JL. 1988b. Sarcomere length and joint kinematics during torque production in frog hindlimb. American Journal of Physiology, 254: C759–68.

Lieber, RL, Shoemaker, SD. 1992. Muscle, joint, and tendon contributions to the torque profile of frog hip joint. American Journal of Medicine, 263(3 Pt 2): R586–R590.

Lieber, RL, Ward, SR. 2011. Skeletal muscle design to meet functional demands. Philosophical Transactions of the Royal Society B: Biological Sciences, 366(1570): 1466–1476.

Lindstedt, SL, LaStayo, PC, Reich, TE. 2001. When Active Muscles Lengthen: Properties and Consequences of Eccentric Contractions. Physiology, 16(6): 256–261.

Llewellyn, ME, Barretto, RPJ, Delp, SL, Schnitzer, MJ, Program, B, Clark, JH, Engineering, B. 2010. Minimally invasive high-speed imaging of sarcomere contractile dynamics in mice and humans. Nature, 454(7205): 784–788.

Lloyd, DG, Besier, TF. 2003. An EMG-driven musculoskeletal model to estimate muscle forces and knee joint moments in vivo. Journal of Biomechanics, 36: 765–776.

Lutz, GJ, Rome, LC. 1996. Muscle function during jumping in frogs. I. Sarcomere length change, EMG pattern, and jumping performance. The American Journal of Physiology, 271: C563–70.

MacIntosh, BR, MacNaughton, MB. 2005. The length dependence of muscle active force: considerations for parallel elastic properties. Journal of Applied Physiology (Bethesda, Md. : 1985), 98(5): 1666–73.

Marsh, RL, Ellerby, DJ, Henry, HT, Rubenson, J. 2006. The energetic costs of trunk and distal-limb loading during walking and running in guinea fowl Numida meleagris I. Organismal metabolism and biomechanics. Journal of Experimental Biology, 209: 2050–2063.

Mayfield, DL, Launikonis, BS, Cresswell, AG, Lichtwark, GA. 2016. Additional in-series compliance reduces muscle force summation and alters the time course of force relaxation during fixed-end contractions. Journal of Experimental Biology, 219(September): jeb.143123.

Morgan, DL, Brockett, CL, Gregory, JE, Proske, U. 2002. The role of the length-tension curve in the control of movement. Sensorimotor Control of Movement and Posture, 508: 489–494.

Rack, P, Westbury, DR. 1969. The effects of length and stimulus rate on tension in the isometric cat soleus muscle. Journal of Physiology, 204: 443–460.

Rack, PM, Westbury, DR, Rack, PM. 1969. The effects of length and stimulus rate on tension in the isometric cat soleus muscle. Journal of Physiology, 204(2): 443–460.

Ramsey, R. W. & Street, SF. 1941. The isometric length-tension diagram of isolated skeletal muscle fibers of the frog. Protoplasma, 36(1): 157–190.

Rassier, DE, MacIntosh, BR, Herzog, W. 1999. Length dependence of active force production in skeletal muscle. Journal of Applied Physiology, 86(5): 1445–1457.

Roszek, B, Baan, GC, Huijing, PA. 1994. Decreasing stimulation frequency-dependent length-force characteristics of rat muscle. Journal of Applied Physiology, 77(4): 2115–2124.

Rubenson, J, Marsh, RL. 2009. Mechanical efficiency of limb swing during walking and running in guinea fowl (Numida meleagris). Journal of Applied Physiology, 106(1985): 1618–1630.

Rubenson, J, Pires, NJ, Loi, HO, Pinniger, GJ, Shannon, DG. 2012. On the ascent: the soleus operating length is conserved to the ascending limb of the force-length curve across gait mechanics in humans. Journal of Experimental Biology, 215(20): 3539–3551.

Rubenson, J, Sanghvi, H, Cromie, MJ, Easton, K, Marsh, RL, Delp, SL. 2013. Influence of tendon compliance and activation level on fibre operating lengths of skeletal muscle. Annual Meeting of the Society for Integrative and Comparative Biology, San Fancisco, CA.

Seth, A, Hicks, JL, Uchida, TK, Habib, A, Dembia, L, et al. 2018. OpenSim : Simulating musculoskeletal dynamics and neuromuscular control to study human and animal movement. Computational Biology, 14(7): e1006223.

Smith, NC, Payne, RC, Jespers, KJ, Wilson, AM. 2007. Muscle moment arms of pelvic limb muscles of the ostrich (Struthio camelus). Journal of Anatomy, 211(3): 313–324.

Sugisaki, N, Kawakami, Y, Kanehisa, H, Fukunaga, T. 2011. Effect of muscle contraction levels on the force-length relationship of the human Achilles tendon during lengthening of the triceps surae muscle-tendon unit. Journal of Biomechanics, 44(11): 2168–2171.

TerKeurs, H, Isazumi, K, Pollack, GH. 1978. The Sarcomere Length-Tension Relation in Skeletal Muscle. Journal of General Physiology, 72(October): 565–592.

Zajac, FE. 1989. Muscle and tendon: properties, models, scaling, and application to biomechanics and motor control. Critical Reviews in Biomedical Engineering, 17(4): 359–411.

