## Supplemental Methods Model Development for "The interaction of compliance and activation on the force-length operating range and force generating capacity of skeletal muscle"

### Supplementary Material: Model Development

#### Glossary

| | |
|---|---|
| Joint Coordinate System | JCS |
| Bone Technical Coordinate Systems | B-TCS |
| Bone Anatomical Coordinate System | B-ACS |
| Global Coordinate System | GCS |
| Optimal fiber length adjusted pennation angle | $\theta_0$ |
| Tendon Slack Length | $L_{ts}$ |
| Average sarcomere length | $L_s$ |
| Average fascicle length | $L_f$ |
| Optimal fiber length | L0 |
| Tarsometatarsus-phalangeal | TMP |

#### Animals

Complete muscle-tendon paths, bone geometry and muscle architecture measurements were made on one guinea fowl specimen (1.45 kg body mass) to construct a generic musculoskeletal model of the pelvic limb. Four additional animals (1.46 ± 0.1 kg; mean ± SD) were used to compare general muscle and tendon properties (muscle mass, fiber length, pennation angle, tendon length and mass). Experimental moment arm measurements were performed on two animals (1.55 kg; 1.49 kg) for ankle and tarsometatarsus-phalangeal (TMP) muscles and on four animals (1.59 ± 0.1 kg; mean ± SD; taken from Carr et al., 2011) for the moment arm of the knee extensors (patella) and hip extensor muscle (the ILPO; see muscle abbreviations Table S1). These experimental moment arms were compared to those predicted by the model. *In vivo* passive joint moment experiments were performed on four animals (1.55 ± 0.2 kg; mean ± SD) and compared to the generic model predictions. Two animals (1.43 kg; 1.49 kg) were used to measure tendon elastic modulus. Animal experiments were performed under protocols approved by the Northeastern University Institutional Animal Care and Use Committee (NU IACUC) and all specimens used only for anatomical measurements were obtained post euthanasia from NU IACUC approved protocols. The model specimen and the additional muscle architecture specimens were transferred to Stanford University Neuromuscular Biomechanical Laboratory for model development.

**3D muscle-tendon paths**

The model animal pelvic limb was skinned and divided down the mid-line of the pelvis. The right limb was kept fresh/frozen while the left limb was fixed in 10% neutral buffered formalin in a posture representing the mid-swing of gait. The three-dimensional (3D) paths of the muscle-tendon units of the fresh/frozen limb were digitized using an optical tracking system (Polaris, Northern Digital, Waterloo, ON) using a protocol similar to that of Hutchinson et al. (Hutchinson et al. 2015) (for a list of muscle-tendon-units incorporated in the model see Table S1). A passive retro-reflective three-marker digitizing probe was used to identify the 3D location of points on the specimen relative to an active LED-emitting marker cluster serving as the Global Coordinate System (GCS) (AdapTrax trackers; Traxal Inc., Toronto, ON). Because the LED-emitting cluster was too large to place directly into the specimen bones, it was rigidly mounted on a frame that could also clamp the specimen securely in place. Before each muscle tendon path was digitized, Bone Technical Coordinate Systems (B-TCS) were identified by digitizing three small non-collinear holes made in each bone (for a flow chart of spatial transformations used in the construction of the musculoskeletal model see Fig. S1). For the pelvis, the B-TCS points were

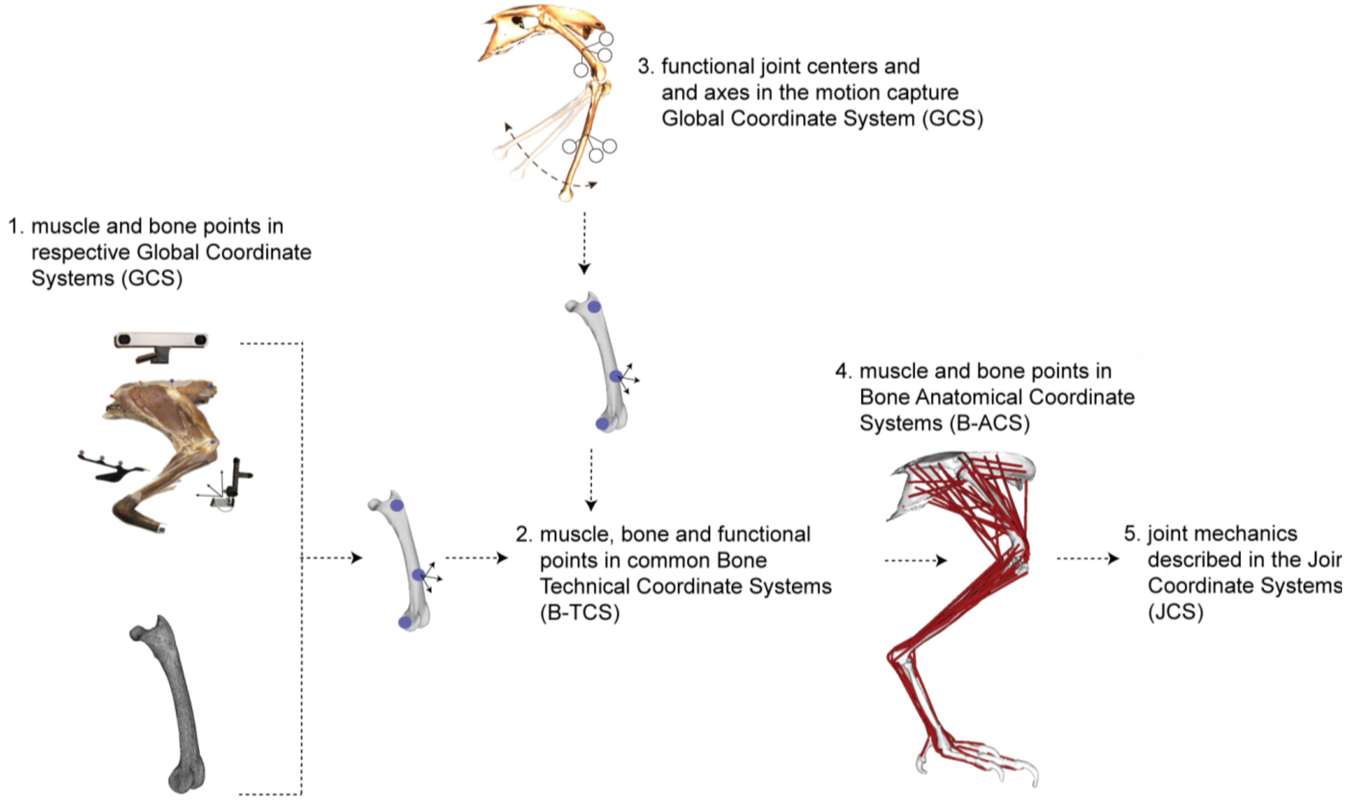


***Figure S1***. *Flow chart for converting bone and muscle points collected in instrument Global Coordinate Systems (GCS) into the Bone Anatomical Coordinate Systems (B-ACS) used to define the Joint Coordinate System (JCS).*

anatomically relevant landmarks that were also used to construct the pelvis Bone Anatomical Coordinate System (B-ACS; see below and Table S2; Fig. S2). B-TCSs were identified for the bones on which the muscle originated and inserted and any bones that the muscle crossed. If the specimen needed to be repositioned in the frame, new B-TCSs points were taken. When tracking the muscle-tendon paths, special care was taken to identify the origin and insertion of the muscle and any anatomical features constraining the muscle-tendon path. After digitizing the muscle-tendon paths, the points relative to the GCS were transformed into the relevant B-TCSs using MATLAB (The MathWorks, Natick, MA). One exception to the use of B-TCSs was for the tendon paths on the phalanges. For these, it was found to be more accurate to define the muscle points based off of the bone geometry directly, guided by careful dissection.

For muscles with broad origins and/or insertions, or for those muscles with a complex architecture (e.g. multi-pennate muscles), we divided the muscle into multiple (2-3) muscle lines of action (see Table S1)[‡]. In these cases, the architecture of the muscle was measures and computed separately for each line of action (see below). Some muscles of the guinea fowl have branched tendons that insert on separate limb segments. For example, the flexor digitorum longus (FDL) muscle inserts on the distal phalanx of digit II, III and IV. For these muscles we generated ‘cloned’ muscles for each tendon branch. The cloned muscles have identical muscle architecture, but different tendon paths.[‡] See Table 1 for details.

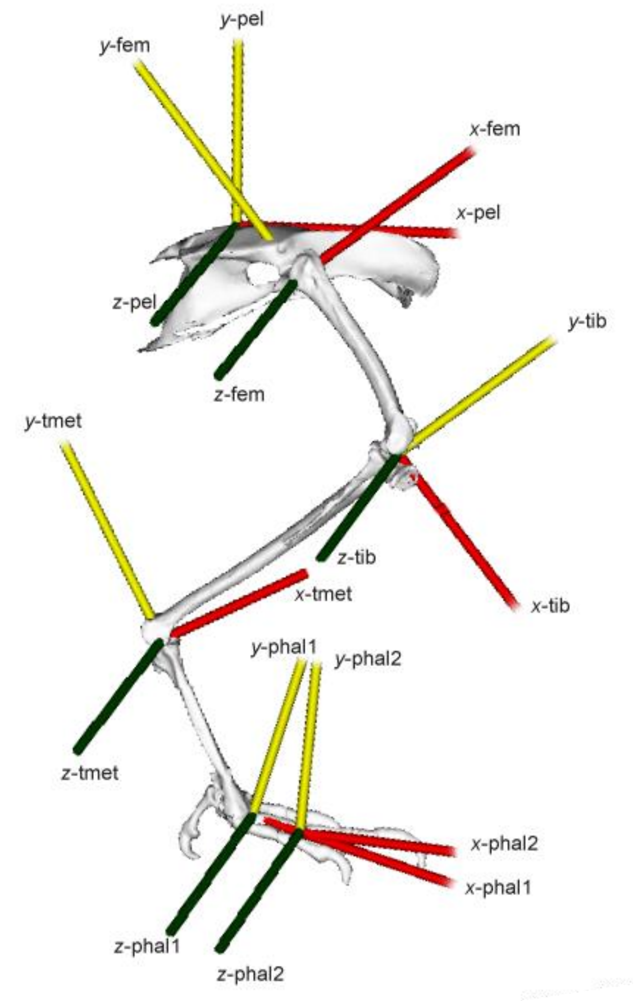


***Figure S2**. Graphical representation of Bone Anatomical Coordinate Systems (B-ACS) for the pelvis (pel); femur (fem); tibiotarsus (tib); tarsometatarsus (tmet); proximal phalanx (phal1); and disatal phalanx (phal2).*

---

[‡] Note that care should be taken when choosing how the multiple lines of action for single muscles and cloned muscles are used in muscle-tendon or joint mechanics simulations. Optimization-based simulations using multiple lines of action for muscles might, in some cases, result in solutions that are not physiologically plausible. For example, vastly different activation patterns/levels across lines of action might not be physiologically feasible if separate neural innervation of the muscle regions (lines of action) are not present. Multiple cloned muscles should typically not be used together since doing so will double the size of the actual muscle. Choosing which cloned muscle to use will depend on the specific modeling performed (for example, which digit kinematics are tracked). For most modelling we recommend using the muscle clone that inserts on digit III since this digit is typically used in kinematic analyses. The data reported in the accompanying manuscript uses the digit III muscle clones. For muscles that insert only on digit II and IV, we recommend constraining digit II and digit III kinematics to the TMP.

***Table S1.*** *Muscle names, abbreviations and description. Muscles with superscript 'C' demotes a 'cloned' muscle for each tendon branch. For 'clone' muscles the underlined muscle is the one used for analysis in the accompanying manuscript. Muscles with superscript 'M' demotes a muscle divided into multiple lines of action. (Some muscles of the guineafowl pelvic limb have been omitted. We do not include the caudofemoralis pars caudalis because we did not model the tail vertebrae. We also do not include the fibularis brevis, a small muscle with a tendon crossing lateral to the ankle, and other small digital extensors originating on the tarsometatarsus).*

| Muscle Name | Abbreviation | Description |
|---|---|---|
| **Ambiens** | AMB | |
| **Caudofemoralis pars pelvica** | CFP | |
| **Extensor Digitorum Longus** [C] | EDL_ii | inserts on digit ii |
| | EDL_iii | inserts on digit iii |
| | EDL_ iv | inserts on digit iv |
| **Femerotibialis intermedius** [M] | FTI_l | lateral |
| | FTI_m | medial |
| **Femerotibialis lateralis pars distalis** | FTLD | |
| **Femerotibialis lateralis pars proximalis** [M] | FTLP_l | lateral |
| | FTLP_m | medial |
| **Femerotibialis medialis** [M] | FTM_p | prox |
| | FTM_m | mid |
| | FTM_d | distal |
| **Fibularis longus** [C] | FL_p | runs posterior to ankle joint |
| | FL_l | runs lateral to ankle joint |
| **Flexor cruris medialis** | FCM | |
| **Flexor cruris lateralis pars accessoria** | FCLP_c | includes the combined pars pelvica and pars accessoria muscles |
| **Flexor cruris lateralis pars pelvica** | FCLP_p | only isolated pars pelvica |
| **Flexor digitorum longus** [C] | FDL_ii | inserts on digit ii |
| | FDL_iii | inserts on digit iii |
| | FDL_iv | inserts on digit iv |
| **Flexor hallucis longus** [C] | FHL_h | inserts on hallux |
| | FHL_ii | inserts on digit ii |
| | FHL_iii | inserts on digit iii |
| | FHL_iv | inserts on digit iv |
| **Flexor perforans et perforatus digiti II** | DFII_s | superficial; crosses knee |
| **Flexor perforans et perforatus digiti III** | DFIII_s | superficial; crosses knee |
| **Flexor perforatus digiti II** [M] | DFII_d | deep; does not cross knee |
| | DFII_dx | deep; crosses knee |
| **Flexor perforatus digiti III** [M] | DFIII_d | deep; does not cross knee |
| | DFIII_dx | deep; crosses knee |
| **Flexor perforatus digiti IV** | DFIV_s | superficial; crosses knee |
| **Gastrocnemius intermedia** | IG | |
| **Gastrocnemius lateralis** | LG | |
| **Gastrocnemius medialis** [M] | MG_l | lateral |
| | MG_c | center |
| | MG_m | medial |
| **Iliofemoralis externus** | IFE | |

| | | |
|---|---|---|
| **Iliofibularis** [M] | IF_p | posterior |
| | IF_a | anterior |
| **Iliotibialis cranialis** [M] | IC_cr | cranial |
| | IC_cd | caudal |
| **Iliotibialis lateralis pars postacetabularis** [M] | ILPO_cr | cranial |
| | ILPO_m | mid |
| | ILPO_cd | caudal |
| **Iliotibialis lateralis pars preacetabularis** [M] | ILPR_cr | cranial |
| | ILPR_cd | caudal |
| **Iliotrochantericus pars caudalis** [M] | ITC_v | ventral |
| | ITC_d | dorsal |
| **Iliotrochantericus pars cranialis** | ITCR | |
| **Iliotrochantericus pars medialis** | ITM | |
| **Ischiofemoralis** [M] | ISF_v | ventral |
| | ISF_d | dorsal |
| **Obturatorius medialis** [M] | OM_v | ventral |
| | OM_d | dorsal |
| **Pubo-ischio-femeralis pars medialis** [M] | PIFM_ cd | caudal |
| | PIFM_cr | cranial |
| **Pubo-ischio-femoralis pars lateralis** [M] | PIFL_ cd | caudal |
| | PIFL_cr | cranial |
| **Tibialis cranialis** [M] | TC_f | muscle head inserts on femur |
| | TC_t | muscle head inserts on tibiotarsus |

#### Bone and Joint Coordinate Systems

Bone Anatomical Coordinate Systems (B-ACS) were generated for each bone from bone landmarks and functional joint centers and axes of rotation. The range of motion of the knee, ankle, and tarsometatarsus-phalageal (TMP; digit III) joints of the fresh/frozen limb were tracked as they were cycled through flexion/extension using a 10-camera motion capture system (Vicon MX100; 200Hz; Oxford Metrics, Oxford, UK). Clusters of three retro-reflective markers were attached to the femur, tibia, tarsometatarsus and the first phalange of digit III using a small bone pin, allowing the relative 3D position and orientation of the adjacent segments to be determined. An average helical flexion/extension axis was computed for the knee, ankle and TMP using the techniques outlined in (Besier et al. 2003) and (Rubenson et al. 2007). The digitizing probe was used to identify the medial and lateral boundaries of the joint and used to establish the functional joint center along the helical axis (see Rubenson et al. 2007). The helical axes and joint centers were initially expressed in the segment cluster coordinate systems. The helical axes and joint centers, together with anatomical landmarks digitized in the motion capture session were used in the construction of the B-ACSs. Digitized landmarks included the synsacrum, sulcus, ilium, pubis and the acetabulum center for the pelvis. Other digitized landmarks included the medial and lateral points of the knee, ankle, TMP and interphalangeal joint (IP, digit III) joints and the medial and lateral points of the end of the third phalange of digit III. The details of how the functional and anatomical coordinates were used in the construction of the B-ACS are outlined in Table S2 and displayed graphically in Figure S2. The B-TCS points were also digitized in the motion capture trials relative to the marker clusters. This step allowed both muscle-tendon paths and bone geometry to be translated from the B-TCSs into the B-ACSs (see Figs. S1&2).

***Table S2****: Description of Bone Anatomical Coordinate Systems (B-ACS).*

| Segment | B-ACS definition | Landmark definitions |
|---|---|---|
| Pelvis | **origin**: SUL<br>**x-axis**: unit vector from SUL to IL (+ cranial)<br>**y-axis**: cross product of x-axis and unit vector from PUB to SYN (+ dorsal)<br>**z-axis**: cross product of x-axis and y-axis | HJC: hip joint center<br>SUL[1]: Sulcus<br>IL[2]: ilium<br>PUB: caudal end of pubis |
| Femur | **origin**: HJC<br>**y-axis**: unit vector from KJC to HJC (+ proximal)<br>**z-axis**: cross product of y-axis and unit vector from MKHA to LKHA (+ lateral)<br>**x-axis**: cross product of y-axis and z-axis. | HJC: hip joint center<br>KJC: knee joint center<br>MKHA: medial knee helical axis endpoint<br>LKHA: lateral knee helical axis endpoint |
| Tibiotarsus | **origin**: KJC<br>**y-axis**: unit vector from AJC to KJC (+ proximal)<br>**z-axis**: cross product of y-axis and unit vector from MKHA to LKHA (+ lateral)<br>**x-axis**: cross product of y-axis and z-axis. | KJC: knee joint center<br>AJC: ankle joint center<br>MKHA: medial knee helical axis endpoint<br>LKHA: lateral knee helical axis endpoint |
| Tarsometatarsus | **origin**: AJC<br>**y-axis**: unit vector from TMP to AJC (+ proximal)<br>**z-axis**: cross product of y-axis and unit vector from MAHA to LAHA (+ lateral)<br>**x-axis**: cross product of y-axis and z-axis. | AJC: ankle joint center<br>TMP: tarsometatarsus-phalangeal joint center<br>MAHA: medial ankle helical axis endpoint<br>LAHA: lateral ankle helical axis endpoint |

| | | |
|---|---|---|
| Proximal Phalanx* | **origin**: TMP<br>**x-axis**: unit vector from TMP to IP (+ cranial)<br>**z-axis**: cross product of y-axis and unit vector from MTHA to LTHA (+ lateral)<br>**x-axis**: cross product of x-axis and z-axis. | TMP: tarsometatarsus-phalangeal joint center<br>IP: inter-phalangeal joint center<br>MAHA: medial ankle helical axis endpoint<br>LAHA: lateral ankle helical axis endpoint |
| Distal Phalanx* | **origin**: IP<br>**x-axis**: unit vector from IP to PHAL (+ cranial)<br>**z-axis**: cross product of y-axis and unit vector from MTHA to LTHA (+ lateral)<br>**x-axis**: cross product of x-axis and z-axis. | IP: inter-phalangeal joint center<br>PHAL: distal end of digit<br>MAHA: medial ankle helical axis endpoint<br>LAHA: lateral ankle helical axis endpoint |

[1]Sulcus; caudal end of the prominent ridge on the midline of the dorsal aspect of the postacetabular ilium.
[2]Ilium; cranial aspect of the ilium, where it meets the sixth thoracic vertebrae
*The B-ACS is the same for digit II, III and IV based on respective TMP and IP joint centers

The musculoskeletal model uses a non-orthogonal Joint Coordinate System convention (JCS) whereby each joint's motion is expressed by three ordered rotations (Grood and Suntay 1983): The first rotation is about the proximal segment's BCS z-axis (flexion/extension rotation); the last rotation is about the distal segment's BCS y-axis (the internal/external rotation rotation); the second rotation is about a floating axis that is perpendicular to the first and last rotation axes (the abduction/adduction rotation). The rotation of the JCS is calculated from the rotation matrix of the distal BCS relative to the proximal BCS. Output muscle and joint moments and joint reaction forces from the model are expressed in the JCSs.

The model included two modifications from the standard JCS construction. First, the model incorporated translation between the tarsometatarsus and tibia BCSs as a function of the ankle flexion/extension rotation. This was necessary to describe the *in vivo* path of the bones during rotation. The bone translation was measured during motion capture trials and implemented in the SIMM joint modeling structure. Secondly, we modeled the patella-knee kinematics as a function of the knee flexion/extension rotation. This was achieved by digitizing the patella location relative to the femur and tibia (B-ACSs) across the joints range of motion. The patella motion was similarly implemented in the SIMM joint modeling structure.

#### Bone Geometry

To visualize the bones in the model we generated high-resolution .ply models of the major leg and foot bones. After digitizing muscle tendon paths and performing the joint motion capture experiments bones of the right leg were de-fleshed and cleaned. The cleaned bones were scanned using a 3D scanner. The pelvis, femur, and tarsometarsus were scanned individually, and the phalangeal segments were scanned together. Bones scans were initially segmented using Mimics software (Materialise, Leuven, Belgium). The phalanges of the foot were separated in software by estimating the location of the center of rotation between adjacent segments. The 3D location of B-TCS points drilled in the bones (the anatomical landmarks making up the B-ACS in the case of the hip) were digitized in Blender software (Blender 2.4; blender.org; Amsterdam, The Netherlands). By knowing the translation matrix of the B-ACS in the B-TCS, the individual vertices of the bone files could be translated into the B-ACS (Figure S1, S2). This was performed in MATLAB. The patella was not individually scanned, so a small disc was modelled to represent the patella in the SIMM model on to which muscles attached.

#### Muscle Architecture

The right limb of the model specimen and other muscle architecture specimens were used to measure muscle and tendon mass and length. The left limbs were formalin-fixed in a mid-swing posture and used for muscle fiber length, sarcomere length and pennation angle measurements.

Muscle and free tendon masses were recorded (nearest 0.1 mg). Free tendon length was measured using a digital caliper. A small bundle of fascicles were dissected free from the fixed muscles and their length was measured (nearest 0.1mm) taking into account fascicle curvature. We performed three fascicle length measurements per muscle (or muscle sub-unit). Sarcomere lengths in each fascicle bundle was measured from second harmonic generation using two-photon laser microendoscopy (see Cromie et al. 2013 for a detailed description of the technique). Briefly, near infrared light at a wavelength of 960 nm (Titanium:Sapphire laser; Chameleon, Coherent, Santa Calra, CA) directed at the fascicle bundles interacted with the myosin-containing A-band of the sarcomeres allowing them to be imaged at high resolution. Sarcomere lengths were calculated from the recorded second harmonic generated images by determining the spatial frequency in the sarcomere pattern's two-dimensional discreet Fourier transform (Cromie et al. 2013) with a custom MATLAB script. Sarcomere lengths were measures over 100 frames of stable images. We measured sarcomere lengths at three locations along the muscle fascicle. The average computed sarcomere lengths ($L_s$) and the average fascicle lengths ($L_f$) were combined to compute the optimal fiber length (L0) using the known optimal sarcomere length (2.36 µm) of guinea fowl muscle (Carr, Ellerby, and Marsh 2011):

$$L_0 = L_f \cdot {}^{2.36}/_{L_s} \qquad \text{(Eq. S1)}$$

Pennation angle of the fixed muscle was measured by first cutting into the muscle to better identify the fascicle orientation relative to the tendon line of action. Pennation angles were measured under magnification using a protractor. The cross-sectional area (*CSA*) of the muscle (or muscle sub-unit) was calculated from the optimal fiber length ($L_0$), muscle mass ($m_{mus}$) and assumed density ($\rho_{mus}$, 0.00106 g/mm$^3$) according to equation S2.

$$CSA = \frac{m_{mus}}{\rho_{mus} \cdot L_0} \quad \text{(Eq. S2)}$$

Maximal isometric force for each muscle (or muscle sub-unit) was calculated using a specific tension of 0.3 N/mm$^2$.

**Tendon properties**

The free common tendon from the lateral, medial and intermedius gastrocnemius muscles (Achilles), and the free tendon from the tibialis cranialis, digital flexor-IV (Flexor perforatus digiti IV) and extensor digitorum longus were used to establish a generic elastic modulus for tendon in the model. Material properties were measured on representative tendons from a combination of two animals using a linear material testing instrument (Bose EnduraTEC, ELectroForce 3200, Framingham, MA, USA). The ends of the tendons were clamped using electronically cooled tissue grips and the tendon section between the clamps was wrapped in cellophane to retain moisture. The displacement of the grips was increased until the tendon first generated measurable force (0.25 N) and then shortened by 0.002 mm to set the tendon segment's slack length ($L_{ts}$). The tendon was programmed to undergo a 5Hz sinusoidal cycle to for a duration of 20s. This cycle frequency resulted in a stretch-shorten cycle that approximated the duration of the stance phase / swing phase of gait. The clamp displacement was programmed such that peak force reached the estimated peak *in vivo* isometric force produced by the muscle (see Muscle Architecture section above) and was established from an initial slow ramped tendon stretch. The tendons displayed creep in force typically over the first 5 cycles. After creep dissipated, 5 stretch-shorten cycles were selected for analysis. Tendon strain was calculated as the displacement of the tendon divided by the tendon slack length. Tendon force was subsequently normalized by the estimated peak isometric force of the muscle. From the resulting strain-normalized force plots we selected data at 10 locations starting at 0 strain and ending at peak strain that captured the shape of the curve (Fig. S3 inset). These were used to set the spline control points in SIMM and OpenSim to predict tendon strain based on simulated muscle force. Muscle-specific spline control points were used for the gastrocnemius muscles, the tibialis cranialis, digital flexor-IV and extensor digitorum longus in the model. We generated a ‘generic’ tendon by averaging the four individual sets of spline control points and used these for all other muscles in the model. Because the tendon strain – force curve was normalized to peak muscle force the tendon properties scale to muscle strength in the model.

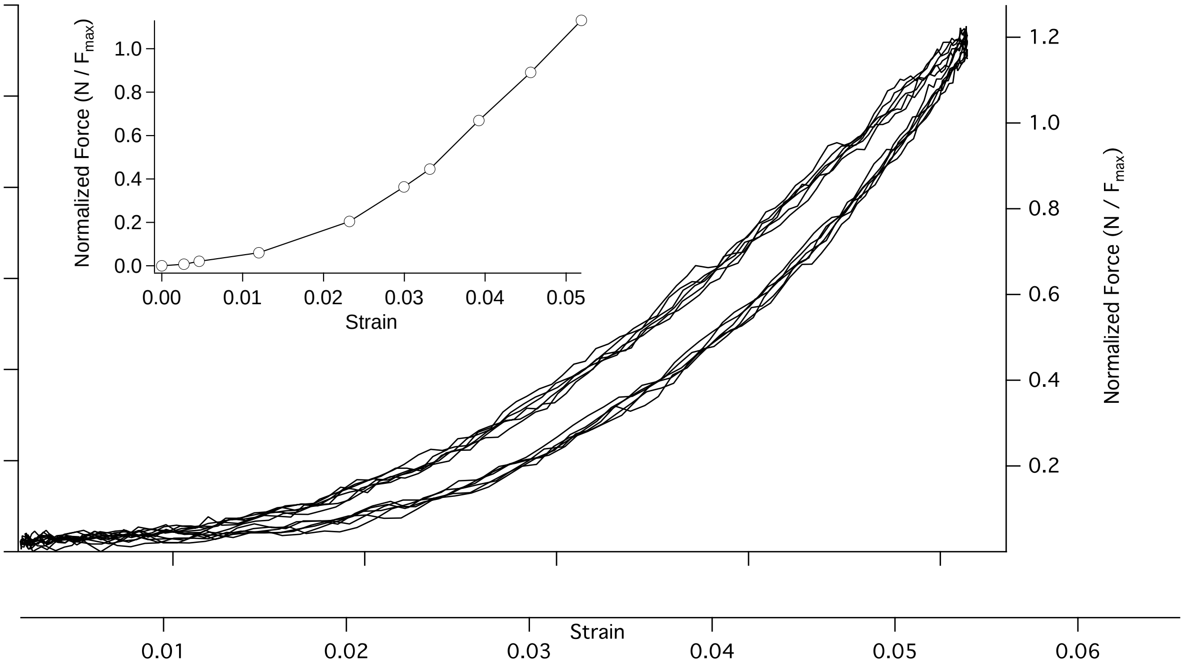


***Figure S3**. Example force-displacement plot of the extensor digitorum longus tendon (5 stretch-shorten cycles). Data is plotted as force vs. displacement and normalized force (force divided by the muscle’s estimated maximum isometric force) vs. strain. The inset illustrates the control points used for the spline fit used in the SIMM and OpenSim modelling environment.*

We also calculated the elastic modulus of the tendons. Tendon stress was calculated as the tendon force (N) divided by its cross sectional area ($CSA_{ten}$; m$^2$). Tendon cross sectional area was computed as:

$$CSA_{ten} = \frac{m_{ten}}{\rho_{ten} \cdot L_{ts}}, \quad \text{(Eq. S3)}$$

where $m_{ten}$ is the mass of the tendon between the clamps (measured to the nearest 0.1 mg) and $\rho_{ten}$ is the density of tendon (0.00112 g/mm$^3$). Elastic modulus was computed as the slope of the ascending linear portion of the stress-strain curve after the tendon toe region. Tendon modulus is reported in Table S2.

| Tendon | Modulus (GPa) |
|---|---|
| **Gastroc. (Achilles)** | 2.48 |
| **Digital Flexor IV** | 1.03 |
| **Tibialis Cranialis** | 0.44 |
| **Extensor digitorum longus** | 0.47 |
| **Generic** | 1.10 |

**Table S2**. Tendon elastic modulus for the individual tendons tested and the average 'generic' tendon.

#### Segment Moment of Inertia

Because we could not empirically measure the segment inertial properties of our model specimen due to muscle dissections, these were predicted from a previous study on guinea fowl joint mechanics (Rubenson and Marsh 2009) which reported the segment mass as a percentage of body mass, and the radius of gyration and location of the center of mass relative to segment lengths. We made a simple conversion of the center of mass location so that it was expressed relative to B-ACS with origins at the proximal joint. The moment of inertial about the ab-adduction axes of the segments (x-axes; y-axis for phalanges) were matched to the flexion-extension values (moment of inertia about the z-axes). The moment of inertia about the long-axis (y-axis) of the femur and other distal segments (x-axis for phalanges) were regarded as small and set to zero. The center of mass location in the segment z-axes (medio-lateral) were also set to zero.

**Model construction in SIMM**

An overview of the model framework is presented in Figure 2 of the accompanying manuscript. Bone geometries and 3D muscle-tendon paths (in relevant B-ACSs) were initially populated in SIMM 6.0 software (Musculographics, Santa Clara, CA). The muscle tendon paths were defined using via points and wrapping surfaces to maintain correct muscle-tendon-unit paths over the joint range of motion. Wrapping surfaces and via points were informed from the experimentally digitized muscle-tendon paths. If required, the origin and insertion points were adjusted slightly so that they resided on the surface of the bone. To minimize discontinuities that result from inaccurate muscle wrapping calculations, muscle paths were edited to constrain the action of wrapping surfaces to between a set of waypoints. In some cases wrapping surfaces were adjusted slightly so that model moment arms were better representative of experimentally measured moment arms (see below).

Muscle optimal fiber lengths (L0), $F_{max}$ (based on *CSA* and a specific tension of 0.3 N/mm$^2$), pennation angle and the tendon force-length curve were input into the SIMM Schutte muscle model. We used the standard SIMM normalized active and passive muscle force length curves (see below for customized activation-dependent active force length curves). Tendon slack length was solved for by constraining the simulated normalized passive muscle fiber length (at the joint posture of the fixed model specimen) to match the experimentally measured normalized fiber lengths. This step took into account the measured pennation angle of the muscle. Custom functions were generated for the ankle and patella kinematics using the SIMM joint editor. Finally, we converted the SIMM version of the model to OpenSim version 3.2 (SimTK, Stanford, CA).

**FL Curve generation**

Activation dependent shifts of the force length curve were implemented in SIMM by creating distinct models with unique force length curves applied to all muscles for each activation level. Activation dependent shifted force length curves were calculated as described below.

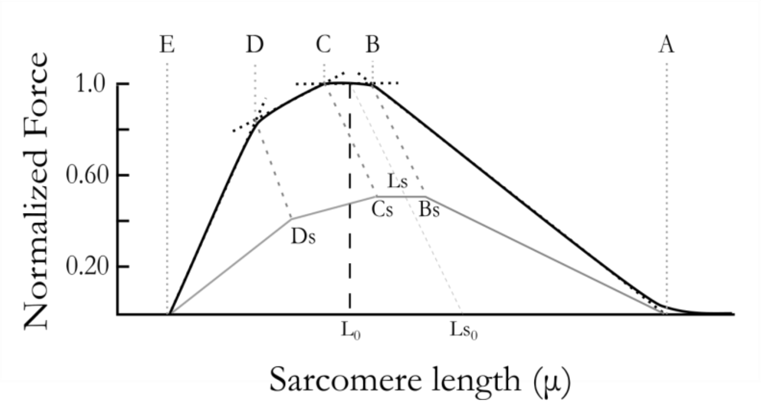


**Figure S4**: Landmarks of the sarcomere length-tension curve (A-E) were shifted as a function of activation (A, Bs, Cs, Ds, E) to generate activation dependent shifted sarcomere length-tension curve. Adapted from Gordon et al. 1966

Starting with previous published values for four limbs of the guinea fowl sarcomere length-tension curve (Carr, Ellerby, and Marsh 2011, E:[1.39,0], C:[2.26,1], B:[2.46,1], A:[3.86,0]), the landmarks of the curve (Gordon, Huxley, and Julian 1966) were shifted with activation level as follows. The length of the plateau was held constant and the relative position of the shift from steep to shallow portion of the ascending limb, Ds, is 60% of the distance between the myosin length, E, and the start of the plateau region, Cs. For two levels of activation-dependent shift (0 and 0.15) and five activation levels (0,0.25,0.50,0.75,1), the x and y coordinates of the landmarks were calculated as:

$$Ls_0 = [L_0*(1+S) , 0]$$

$$Ls = [L_0*(1+S)*A , A]$$

$$Cs = [Ls_x-(B_x-C_x)/2 , A]$$

$$Bs = [Ls_x+(B_x-C_x)/2 , A]$$

$$Ds = [(Cs_x-E_x)*0.6 + E_x, Ls_y*0.8)$$

where $Ls_0$ is the shifted optimal fiber length at no activation, Ls is the shifted optimal fiber length, Cs and Bs the start and end of the plateau region (Fig. S4).

To account for non-uniform striation spacing, variability was added into the length-tension curve by adding +- 0.05 jitter to x and y coordinates of points over 1000 iterations. These data were fit with a $5^{th}$ order polynomial with 23 nodes and the resulting curve was normalized by the original L0. For each activation level, a new SIMM model was created with a new normalized force-length curve as described above.

#### Experimental moment arm measurement

To compare the model-generated muscle moment arms to experimental moment arms we performed tendon travel experiments for muscles from two representative anatomical specimens for the ankle and TMP muscles, and from four animals for the patella and hip muscle (ILPO) moment arms (see ‘Animals’ section above). These comparisons were used to check whether the model’s moment arms arising from input muscle-tendon paths and bone wrapping surfaces were representative (Figs. S5&6). In the case of the gastrocnemius tendon and tibialis cranialis tendon, we adjusted the bone wrapping surfaces in SIMM so that the shape of the moment-arm ankle angle curve provided a closer match to the experimental data. The methods for measuring

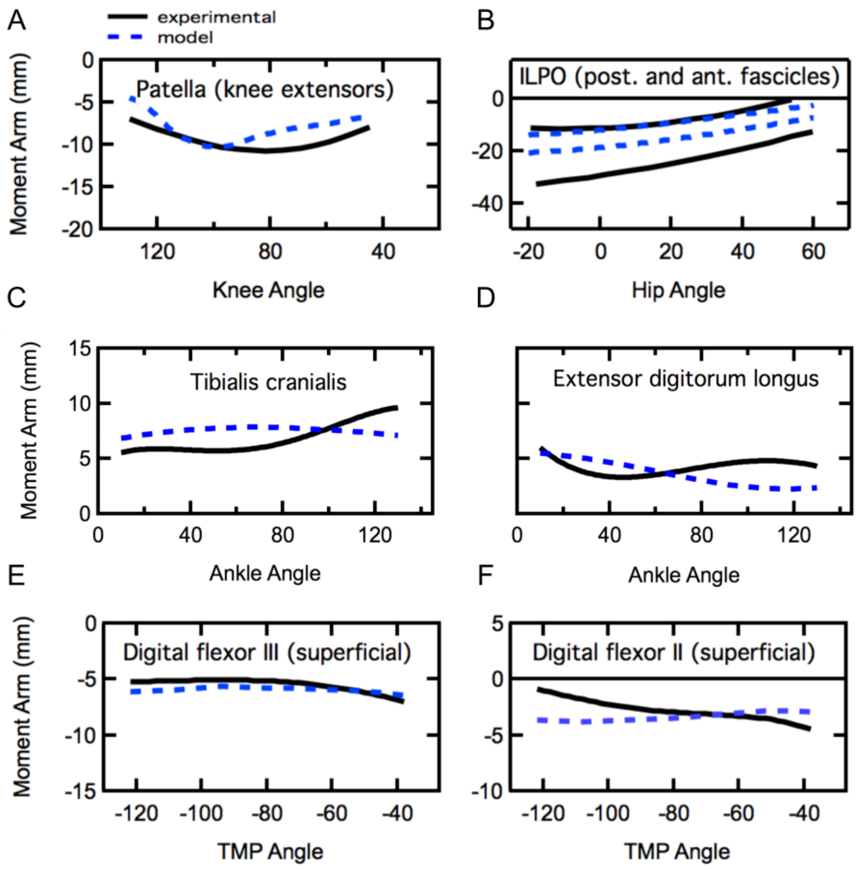


***Figure S5*** *Comparison of model generated moment arms and experimental moment arms for the patella (A), the anterior and posterior fascicles of the ILPO muscle (B), the ankle flexor muscles (C&D) and the TMP flexor muscles (E&F). Note, for the Digital flexor II muscle the moment arm about the TMP was computed from muscle length changes and joint angles while rotating the digit III.*

tendon travel-based moment arms on guinea fowl muscle have been described previously (Carr, Ellerby, and Marsh 2011). Briefly, we combined simultaneous recordings of tendon length (Harvard Bioscience length transducer; Model 52-9511, Holliston, MA, USA; 1000 Hz) and joint angle (digital video; JVC model #GR-DVL9800; JVC, Wayne, NJ, USA; 60 Hz) as the ankle, knee or hip was rotated through its range of motion. The anatomical specimen and limb segments were kept stationary using bone clamps. For hip muscles, we clamped the femur and moved the pelvis segment; for knee muscles (patella tendon moment arm) we clamped the femur and moved the tibiotarsus; for ankle muscles we clamped the tibiotarsus and moved the tarsometatarsus. The individual muscles for which moment arms were measured are shown in Figures S5&6. For the gastrocnemius and digital flexor muscles the muscle tendon unit was left largely intact, with the origin of the muscles separated from the femur or tibiotarsus. These muscles were attached to the length transducer using silk suture with the proximal path of the muscle fibers constrained by a guide glued to the tibiotarsus and/or femur. To measure the moment arm of the patella tendon most of the muscle knee extensor muscle was dissected off but the joint capsules, articulating tissues and small muscles overlying the joints were left intact. The patella tendon was attached via a guide that maintained the *in vivo* path of the tendon over the articulating surfaces of the knee (Carr, Ellerby, and Marsh 2011). The proximal path of the suture was also constrained by guides attached to the femur to replicate the natural path of the knee extensor muscles. For the ILPO moment arm at the hip, the anterior and posterior fascicle paths were identified, after which the muscle was dissected off the pelvis. The fascicle paths were replicated using silk suture that was anchored at the pelvis origin and passed through guides glued to the femur to the length transducer. For all muscles, the length transducer lever was counterweighted to ensure that there was no slack in the suture and that any small strain in

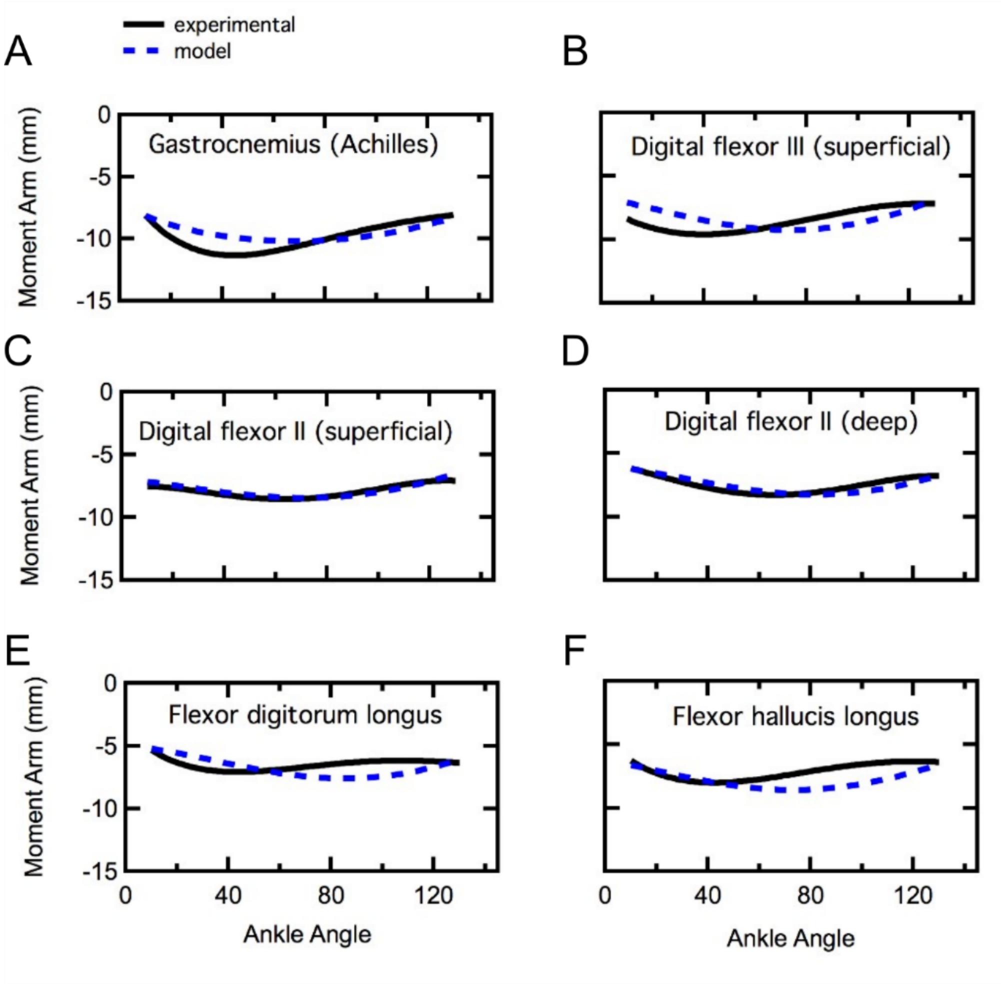


***Figure S6****. Comparison of model generated moment arms and experimental moment arms for the ankle extensor muscles.*

tendon (for the ankle muscles) or suture was kept constant.  The change in length of the muscle-tendon path was recorded as the joint was rotated manually through its range of motion.

Planar joint angles were estimated by digitizing reflective markers placed cranially and caudally on the pelvis and proximally and distally on the femur, tibiotarsus and tarsometatarsus (see Rubenson and Marsh 2009).  The joint center locations were also identified with a reflective marker.   Video recordings were synchronized with length data using a TTL pulse that turned on an LED in the video field of view.  Video data was auto digitized using the Mtrack2 plugin in

ImageJ (ImageJ; NIH, Wayne Rasband; http://rsb.info.nih.gov/nih-image/). Digitized video data was filtered and joint angles were computed using custom MATLAB scripts (MathWorks, Natick, MA, USA). The corresponding length data was filtered and down sampled to match the video data. The length-angle data were fitted with polynomials, the order of which were determined statistically (Carr, Ellerby, and Marsh 2011). These polynomials were subsequently analytically differentiated to yield a moment arm-joint angle equation for each muscle.

**Experimental passive joint moment measurement**

To further assess the accuracy of our model, we compared simulated net passive joint moments to experimental values. Passive joint moments were measured in four animals (see 'Animals' section above) for the hip joint (proximal muscles) and ankle joint (distal muscles). Animals were deeply anesthetized (isoflurane, 1.5%) and core temperature maintained using a heating pad and warm-water sachets placed around the animal. Because the stretch reflex could influence joint moment recordings we also used a local nerve block (Bupivacaine, 0.5%; 5ml) to the pelvic limb nerve supply including the ischiadicus (sciatic) and femoralis nerves. We used a custom limb immobilization rig with the animal positioned on its side that allowed us to freely rotate the joint of interest while immobilizing the adjacent joints at set angles (Fig. S7). This was achieved by securing sliding aluminum brackets across the knee, ankle and TMP joints at specific joint angles using a turn-screw. The brackets were attached directly to the femur, tibia and tarsometatarsus using stainless steel screws secured into pre-drilled holes. The phalanges were positioned using a cable tie secured to the bracket. The hip joint was immobilized using a stage with plastic stops that secured the pelvis and femur in place.

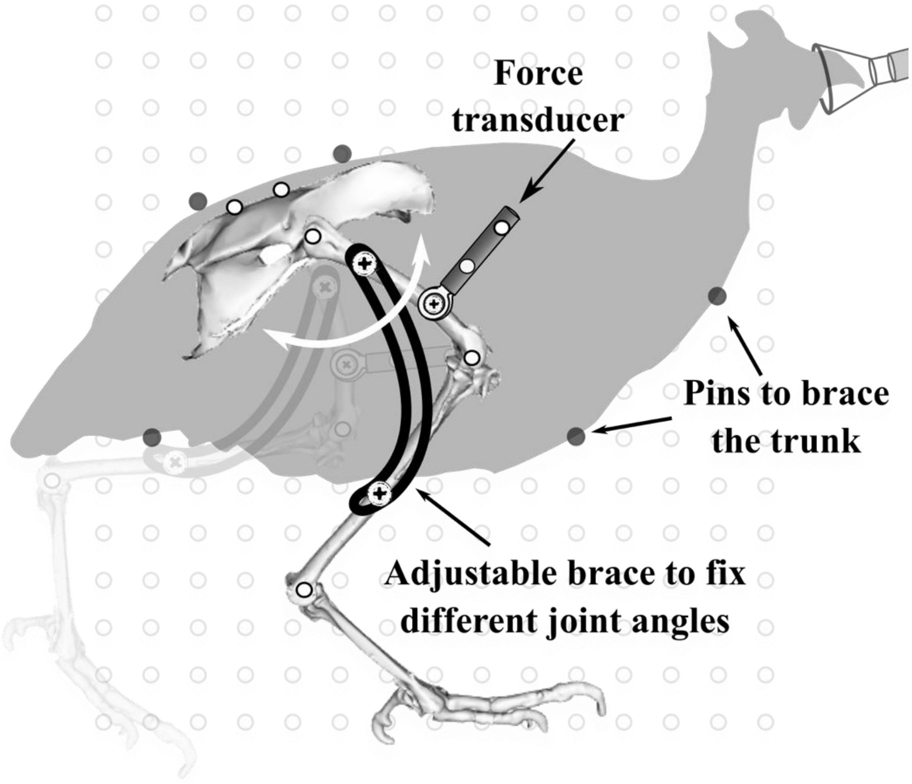


***Figure S7***. *Experimental set-up for passive net joint moment measurements. Example shown for passive hip joint measurement. Also performed for ankle and TMP joints*

We attached a single-axis compression/tension quartz force transducer (Kistler model 9203) to the bone distal to the joint of interest using a stainless-steel mounting screw. The transducer was positioned such that it was held horizontal and so that force applied to the bone segment through the transducer was oriented parallel to the sagittal plane. The transducer was allowed to rotate about its attachment point but was constrained to the sagittal plane (Fig. S7). The joint of interest was held horizontal and rotated through its flexion/extension range of motion by pushing/pulling the force transducer while preventing any other point of force application. Force was recorded continuously (1000 Hz) using a USB A-to-D system (PowerLab, ADInstruments; Bella Vista, Australia). The origin and orientation of the applied force transducer was identified by video recording (60 Hz) a reflective marker positioned at the transducer attachment point and two markers that defined the transducer axis (Fig. S7). The skeletal planar kinematics were recorded from reflective markers placed on joint centers and bone landmarks following the procedures published earlier (Rubenson and Marsh 2009). The force transducer and joint kinematics were synchronized using a TTL pulse that was recorded on a separate A-to-D channel and that simultaneously turned on an LED in the video field of view.

We performed passive joint moment experiments over the joints' range of motion with the adjacent joints set at flexed and extended positions (Figs. S9&10). Joint rotations were performed slowly to minimize acceleration-effects. Pilot experiments confirmed that joint rotation velocity did not greatly alter the passive joint moment profiles. We used an inverse dynamic approach to compute the net joint moments. Similar to moment arm calculations (see above), video data was auto digitized using the Mtrack2 plugin in ImageJ (ImageJ; NIH, Wayne Rasband; http://rsb.info.nih.gov/nih-image/) and synchronized with force data using a TTL pulse that turned on an LED in the video field of view. Digitized video data and force data were filtered (10 Hz Butterworth filter, MATLAB, The MathWorks, Natick, MA, USA). Joint angles and moments were computed similar to the procedure outlined in Rubenson and Marsh (2009) with two modifications to the inverse dynamic model. First, we implemented the external force from the force transducer at the point of attachment to the bone. Second, because the limb and force transducer were held horizontal, the force due to gravity was omitted from our calculations. Moment measurements from individual animals were normalized to body mass. Mean moments and standard deviation of the mean were computed from the four animals over the range of joint angles that were common to all animals. Comparisons of experimental and modelled net passive joint moments for the ankle and hip are presented below in Figures S9&10.

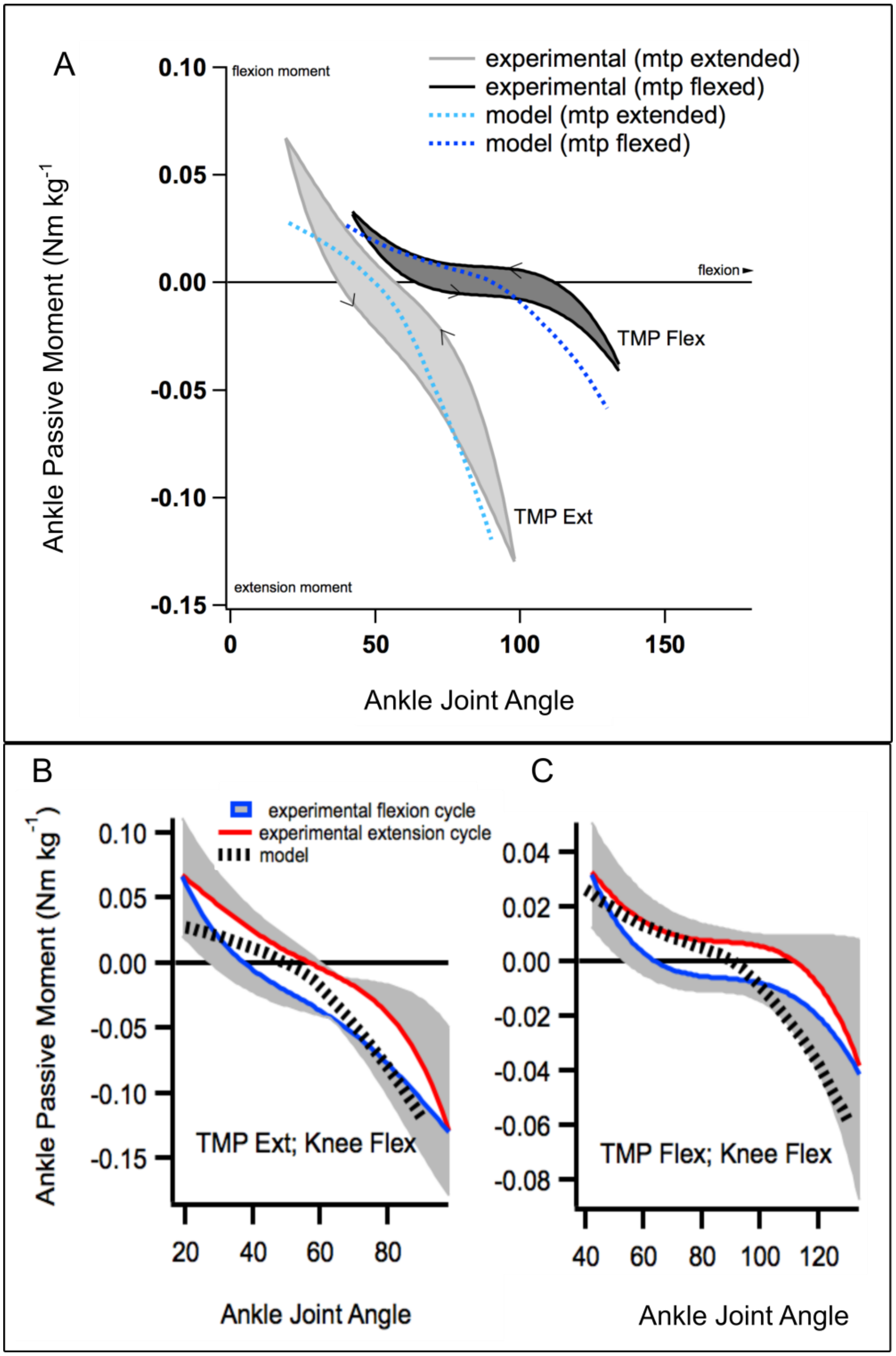


***Figure S9***. *A) Mean passive moments about the ankle as a function of ankle angle, with the TMP joint set at a flexed position (stretching the digital extensor / ankle flexor muscles) and at an extended position (stretching the digital flexor/ ankle extensor muscles). The arrows on the moment curves represent the direction of joint rotation (the shaded region between flexion and extension cycles are included for visual clarity). Modeled moments are overlain for comparison. B&C) Mean passive moment data for the flexion and extension cycles including the standard deviation of the mean (grey shaded regions). Modeled moments are overlain for comparison.*

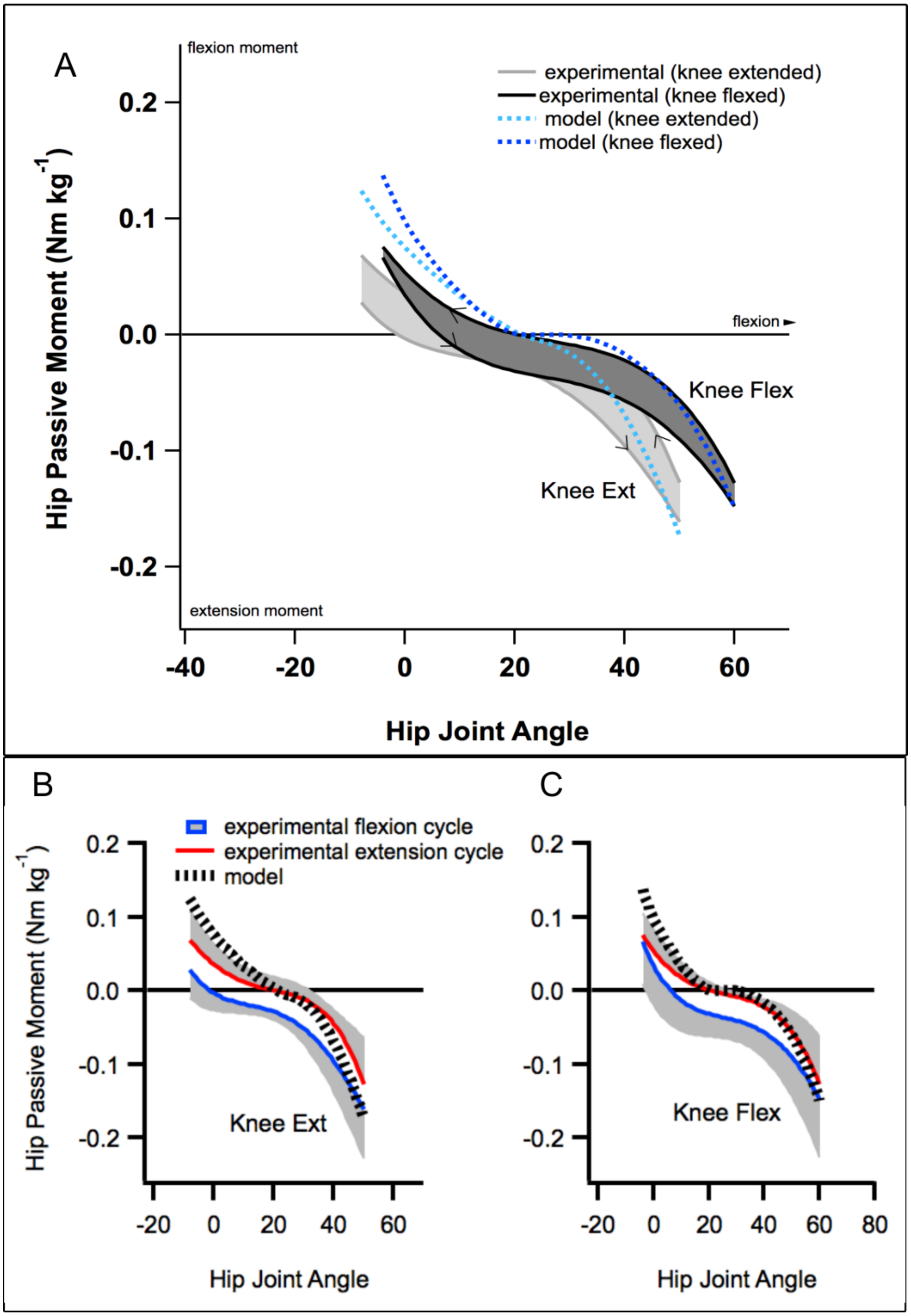


***Figure S10***. *A) Mean passive moments about the hip as a function of hip angle, with the knee joint set at a flexed position (stretching the hip flexor muscles) and at an extended position (stretching the hip extensor muscles). The arrows on the moment curves represent the direction of joint rotation (the shaded region between flexion and extension cycles are included for visual clarity). Modeled moments are overlain for comparison. B&C) Mean passive moment data for the flexion and extension cycles including the standard deviation of the mean (grey shaded regions). Modeled moments are overlain for comparison.*

**References**

Besier, Thor F, Daina L Sturnieks, Jacque A Alderson, and David G Lloyd. 2003. “Repeatability of Gait Data Using a Functional Hip Joint Centre and a Mean Helical Knee Axis.” *Journal of Biomechanics* 36: 1159–68. https://doi.org/10.1016/S0021-9290(03)00087-3.

Carr, Jennifer A, David J Ellerby, and Richard L Marsh. 2011. “Differential Segmental Strain during Active Lengthening in a Large Biarticular Thigh Muscle during Running.” *Journal of Experimental Biology* 214 (Pt 20): 3386–95. https://doi.org/10.1242/jeb.050252.

Cromie, Melinda J., Gabriel N. Sanchez, Mark J. Schnitzer, and Scott L. Delp. 2013. “Sarcomere Lengths in Human Extensor Carpi Radialis Brevis Measured by Microendoscopy.” *Muscle and Nerve* 48 (2): 286–92. https://doi.org/10.1002/mus.23760.

Gordon, a. M., a. F. Huxley, and F. J. Julian. 1966. “The Variation in Isometric Tension with Sarcomere Length in Vertebrate Muscle Fibres.” *The Journal of Physiology* 184 (1): 170–92. https://doi.org/5921536.

Grood, E S, and W J Suntay. 1983. “A Joint Coordinate System for the Clinical Description of Three- Dimensional Motions : Application to the Knee 1.” *Journal of Biomechanical Engineering* 105: 136–44.

Hutchinson, John R, Jeffery W Rankin, Jonas Rubenson, Kate H Rosenbluth, Robert A Siston, and Scott L Delp. 2015. “Musculoskeletal Modelling of an Ostrich (Struthio Camelus) Pelvic Limb: Influence of Limb Orientation on Muscular Capacity during Locomotion.” *PeerJ* 3: e1001. https://doi.org/10.7717/peerj.1001.

Rubenson, Jonas, David G Lloyd, Thor F Besier, Denham B Heliams, and Paul A Fournier. 2007. “Running in Ostriches ( Struthio Camelus ): Three-Dimensional Joint Axes

Alignment and Joint Kinematics.” *Journal of Experimental Biology* 210: 2548–62. https://doi.org/10.1242/jeb.02792.

Rubenson, Jonas, and Richard L Marsh. 2009. “Mechanical Efficiency of Limb Swing during Walking and Running in Guinea Fowl ( Numida Meleagris ).” *Journal of Applied Physiology* 106 (1985): 1618–30. https://doi.org/10.1152/japplphysiol.91115.2008.
